# Single-cell transcriptional changes of artemisinin-sensitive K13^C580^ and artemisinin-resistant K13^580Y^ *Plasmodium falciparum* upon dihydroartemisinin exposure

**DOI:** 10.1101/2023.12.06.570387

**Authors:** Sean V. Connelly, Clark Cunningham, Ian Petrow, Nazrin Rustamzade, Jenna Zuromski, Deborah M. Sipolski, Christian P. Nixon, Jonathan D. Kurtis, Jonathan J. Juliano, Jeffrey A. Bailey, Cliff I. Oduor

## Abstract

Artemisinin-based therapies have been central to malaria control, but partial resistance in *Plasmodium falciparum*, driven by mutations in the Kelch13 (K13) protein, threatens these gains. To investigate the molecular basis of this resistance, we applied single-cell RNA sequencing to coisogenic parasite lines, K13 wild-type (K13^C580^) and the artemisinin-resistant mutant (K13^580Y^), following a 6 hour pulse of dihydroartemisinin (DHA). This approach enabled high-resolution profiling across intraerythrocytic stages. Both lines exhibited stage-specific transcriptional responses, with pronounced changes in ring and trophozoite stages. Using Manifold Enhancement of Latent Dimensions (MELD), a computational framework for quantifying transcriptional perturbation, DHA-treatment induces stage-specific differences in protein export and metabolic pathways in K13^C580^ and K13^580Y^ parasites, relating to an altered metabolic stress response state. GARP, a potential therapeutic target, was highly differentially expressed in untreated ring stages of K13^580Y^ and K13^C580^. Functional assays confirmed that anti-GARP antibodies retained efficacy against K13^580Y^, supporting its potential as a therapeutic target. These findings provide a comprehensive view of the cellular responses related to artemisinin resistance, identify molecular features of pathogenesis, and highlight surface proteins like GARP as promising intervention targets. This work underscores the power of single-cell approaches to dissect drug responses and guide strategies to overcome resistant parasites.

## INTRODUCTION

Malaria, caused mainly by the species *Plasmodium falciparum*, caused an estimated 610,000 deaths in 2024 (1). The World Health Organization recommends artemisinin (ART) based combination therapies (ACTs) to treat uncomplicated falciparum malaria. ACTs rely on the fast-acting properties of the ART derivative to rapidly clear parasites, while the longer-acting partner drug clears residual parasites. This combination reduces the frequent observation of recrudescent infections when ART derivatives are used alone (2). Parasites have evolved ART partial resistance (ART-R), which first emerged in western Cambodia and spread throughout the Greater Mekong Subregion in Southeast Asia (3,4). ART-R has now emerged in Africa and is spreading through the continent, endangering ACT efficacy (5–15). ART-R manifests as a decreased susceptibility of ring-stage parasites to ART. Clinical ART-R is defined as a reduced rate of parasite clearance (>5 hr parasite half life), residual parasitemia 3 days after treatment, or increase rate of recrudescence (16). Increased parasite survival due to ART-R risks the development of partner drug resistance (17). Given Africa suffers the vast majority of disease burden, a public health disaster could arise if ART-R spreads across the continent (18).

ART-R is due to nonsynonymous mutations in the ß-propeller domain of the Kelch13 (K13) protein (6,19,20). The identification of *K13* (*PF3D7_1343700*) gene mutations as a marker of resistance has provided the impetus for world-wide surveillance to catalog, track and analyze the distribution of *K13* mutations in malaria-endemic regions. Over 100 different *K13* mutations have been identified (21,22), with thirteen considered validated resistance markers and ten considered candidate resistance markers (23). The K13^580Y^ mutation predominates in Southeast Asia (4). In Eastern Africa, the mutations K13^561H,675V,469Y,622I^ have emerged and are now found in multiple countries (24). Validated *K13* mutants exhibit about tenfold higher residual viability than their genetically matched wild-type parasites when exposed to dihydroartemisinin (DHA) at the early ring stage (20).

A number of studies have examined the complex molecular response to understand the underpinnings of survival. Different *K13* mutations have been investigated through gene editing and protein localization experiments (20,25–27). While most studies focus on the treatment response of the early ring stage, few studies have focused on the effect of ART treatment over asexual stages *in vitro.* A study of stage specific artemisinin response correlated fluctuations in drug sensitivity to hemoglobin digestion (28). Further, stage specific gene expression changes were identified in artemisinin resistant isolates from Southeast Asia, with rings and trophozoites showing reduced expression of metabolic pathways and schizonts showing increased expression of protein metabolism pathways (29). Single cell RNA sequencing (scRNAseq) allows for the investigation of stage specific gene expression dynamics in ART-R and sensitive lines. scRNAseq has been utilized in multiple cell atlas efforts across malaria species (30,31), but its utility to study gene expression response to treatment perturbations has not been fully harnessed (32).

In this study, we explored the single cell transcriptomic profile of *P. falciparum* coisogenic lines K13^580Y^ (MRA-1251) and K13^C580^ (MRA-1250) following DHA exposure using a well-based single cell capture platform, SeqWell (33–35). We focused on the stage specific transcriptional effects of DHA treatment on the ring and trophozoite *Plasmodium* cell cycle stages. We observed parasite stage specific gene expression profiles and also potential vulnerabilities of the parasite’s response to DHA treatment, such as elevated GARP surface protein expression, that can be a logical therapeutic target for effective combination therapies in the face of ART-R.

## METHODS

### Parasites lines and culture

The strains of parasite lines used in this study were MRA-1250 (K13^C580^), an artemisinin susceptible fast-clearing Cambodian *P. falciparum* isolate, and MRA-1251 (K13^580Y^), a Cambodian *P. falciparum* isolate which features a single nucleotide substitution leading to a C580Y amino acid change (MR4, BEI Resources, Manassas, VA). The *P. falciparum* parasites were maintained at 3% hematocrit in human O+ RBCs and *P. falciparum* culture media comprising RPMI 1640 (Thermo Fisher Scientific) supplemented with 0.5% (w/v) Albumax II, 50 mg/L hypoxanthine, 0.225% NaHCO3, 25mM HEPES, and 10 mg/L gentamycin. Parasites were cultured at 37 °C in 5% O2, 5% CO2, and 90% N2 (36). The parasite lines were cultured for 48hrs before exposing them to DHA and DMSO.

### Drug susceptibility assays

DHA stocks were made in dimethyl sulfoxide (DMSO) and aliquots were stored at −20 °C. All drug assays were conducted such that the final DMSO concentration was <0.5%. Each parasite line was exposed to 700 nM DHA and control vehicle-treated 0.05% DMSO in independent experiments for 6hrs (**Figure 1A**). Samples from each treated condition were collected at 2hr intervals for scRNAseq library preparation.

**Figure 1:**
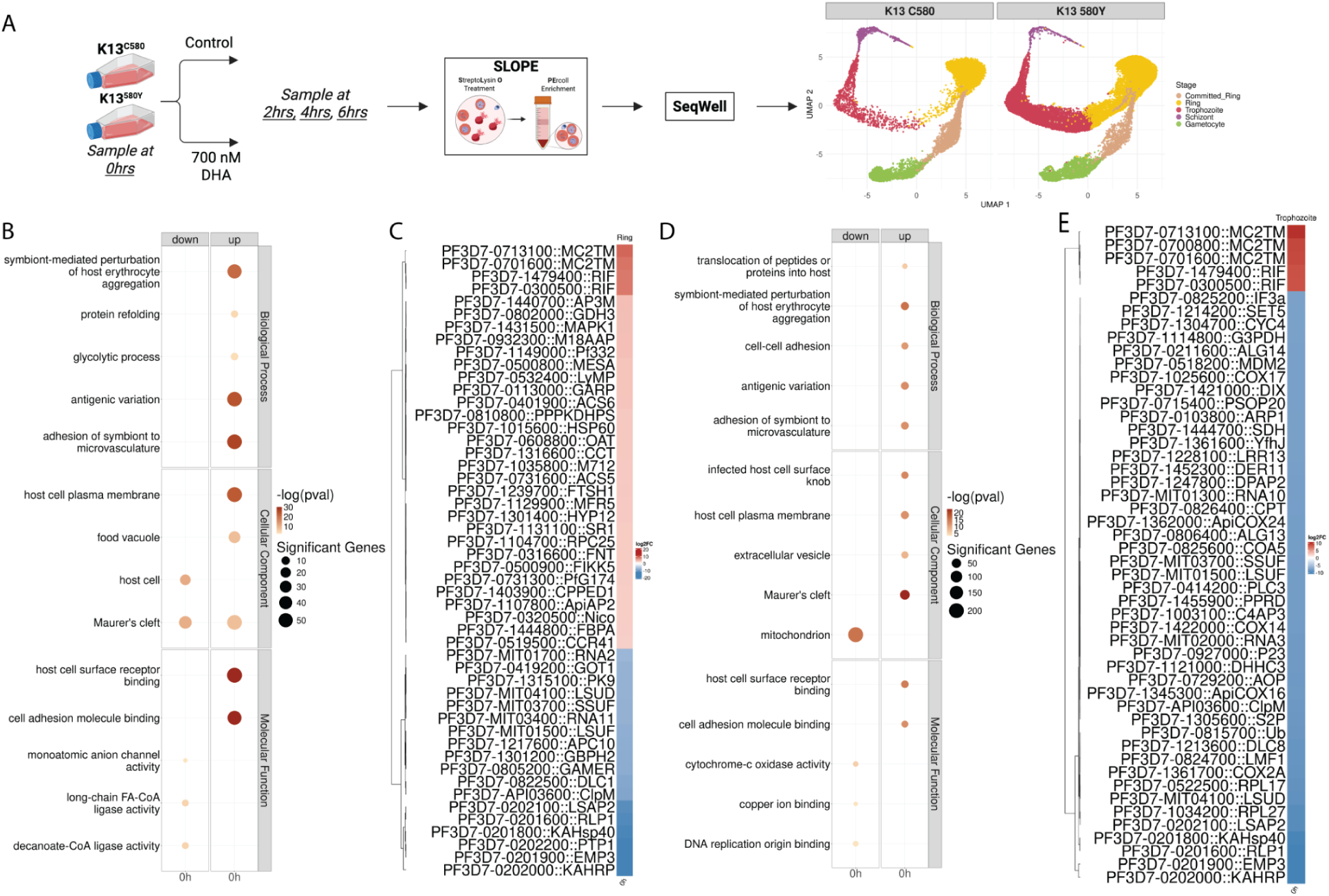
K13^580Y^ vs. K13^C580^ Rings and Trophozoites Showed Upregulation of Parasite-Erythrocyte Interaction Pathways, with Downregulation of Mitochondrial and Apicoplast Genes. **(A)** K13^C580^ and K13^580Y^ cell cultures were grown over 48 hours. A baseline control sample (<u>0h</u>) was collected before splitting each flask into vehicle control (DMSO) and treatment samples (700nM DHA). Over three timepoints (<u>2h</u>, <u>4h</u> and <u>6h</u>) during DHA treatment, we collected samples to enrich via SLOPE. After enrichment, cells were seeded into the SeqWell array for single cell capture and downstream processing. UMAP embedding of the final malaria stage annotation over all datasets is shown, colored by stage (Brown: Sexually Committed Ring, Yellow: Ring, Red: Trophozoite, Purple: Schizont, Green: Gametocyte). The graph is separated by strain (left: K13^C580^, right: K13^580Y^). The top 5 significant Gene Ontology (GO) enrichment terms (topGO p<0.01) in each category between K13^580Y^ vs. K13^C580^ rings **(B)** and trophozoites **(D)** are plotted, with the number of significant genes that support that GO Term being the size of the circles and the color being the magnitude of the negative log of the topGO p-value. A heatmap of the 50 differentially expressed (DE) annotated genes by absolute log2FC between K13^580Y^ vs. K13^C580^ rings **(C)** and trophozoites **(E)** are plotted. The top 5 hypervariable genes by absolute log2FC were selected for visualization for each heatmap. An asterisk indicates genes that were significantly differentially expressed at an absolute log2FC > 1 and a BH-adjusted p-value < 0.001. Red boxes are upregulated genes, white boxes indicate genes that were not differentially expressed, and blue boxes are downregulated genes.

### Enrichment of parasite infected RBCs

To prepare the scRNAseq libraries, we enriched the parasite infected RBCs thus increasing chances of loading and capturing mRNA transcripts from the parasites. Our unsynchronized parasite cultures were enriched using SLOPE as previously described (37) with some modifications. Briefly, erythrocyte density was measured using a Cellometer Auto T4 (Nexcelom Biosciences, Lawrence, MA) and cell density adjusted to 2×10^9^ erythrocytes/mL. The 22.5U of the lytic agent streptolysin-O (SLO) per 50µl RBCs was added in a ratio of 2 parts SLO solution to 5 parts erythrocytes. Samples were mixed well by pipetting and incubated at room temperature for 6 min. 10 volumes of 1X PBS or media (RPMI 1640 HEPES) were added and cells were centrifuged at 2,500xg for 3 min. After removal of the supernatant, cells were washed twice more with 1X PBS. Following SLO lysis, cells suspended in 1X PBS were layered onto a 60% Percoll gradient (Sigma Aldrich, St Louis, MO) and centrifuged at 1,500xg for 10 min. The top layer of Percoll was discarded while the lower cell pellet, rich in infected erythrocytes, was collected and washed twice with 1xPBS and later suspended in 200µl RPMI to be loaded into the SeqWell arrays.

### Single-Cell library preparation and sequencing

To capture single cell transcriptomes, we utilized SeqWell, a massively parallel, low-input scRNAseq platform (33–35). Details of array loading, library preparation, and sequencing are provided in the **Supplemental Methods**. Raw read data and processed UMI count matrices from K13^C580^ and K13^580Y^ scRNAseq experiments have been deposited and are publicly available through the SRA database (BioProject ID: PRJNA1049964).

### Single-Cell Transcriptome Analysis

Raw reads were processed using version 3.0.2 of the Drop-seq pipeline (38), and according to the ‘Drop-seq Alignment Cookbook’, both found at http://mccarrolllab.com/dropseq/. Demultiplexed FASTQs were aligned to a concatenated human (Gencode v48) and *P. falciparum* 3D7 reference genome (PlasmoDB v68) using the STAR aligner. Any reads aligned to human chromosomes were filtered out with a custom script (https://github.com/sconnelly007/K13_mt_v_wt_sc/scripts/processing/01_read_filter_human.py). Individual reads were tagged with a 12bp barcode and 9bp unique molecular identifier (UMI) contained in Read 1 of each sequencing fragment. Following alignment, reads were grouped by the 12bp cell barcodes and subsequently collapsed by the 9bp UMI to generate a gene expression count matrix. Gene expression scRNAseq count matrices from K13^C580^ and K13^580Y^ cell line experiments (n = 7 per *Pf* cell line) were analyzed and visualized in R (v4.5.2) using the Seurat package (v5.3.1) (39).

### Quality Control Filtering

During quality control filtering, cells with <300 genes, <500 UMIs and >20% mitochondrial gene expression were removed (**Supplemental Figure S1** and **S2**). Genes were retained if they were expressed in at least 10 cells or expressed in at least 1% of all cells (**Supplemental Figure S3** and **S4**).

### Cell Annotation Pipeline

The annotation strategy is outlined in **Supplemental Methods** and **Supplemental Figure S5**. Each individual sample underwent doublet detection with scrublet (v0.2.3) and doubletFinder (v2.0.6) (**Supplemental Figure S6** and **S7**). The expected percentage of doublets was estimated to be 2.37% based on previously published SeqWell experiments (34). In scrublet, a doublet detection threshold of 0.13 was used over all datasets. In doubletFinder, a per dataset threshold was set using the strategy outlined in the Demuxafy package (40).

**Pseudotime** Pseudotime values and weights were calculated using slingshot over the entire dataset (41) (**Supplemental Figure S14**).

### Fine Grained Differential Expression with MELD

Applying the MELD algorithm and performing differential expression (DE) on our data is detailed in the **Supplemental Methods.** All DE results across stages are in **Supplemental Table S1**.

### Gene Ontology (GO) Analysis

Using the significant differentially expressed genes (DEGs), GO analysis was performed with pfGO (v2.1, https://github.com/oberstal/pfGO), a R wrapper for topGO (42). GO term enrichment was performed similarly to previous studies (43), with the output DE results from MAST filtered to DEGs with BH-adjusted p values < 0.001 and categorized as upregulated (log2FC > 1) or downregulated (log2FC < -1). For each timepoint and in each category, GO term enrichment was performed using a Fisher test with the topGO weight01 algorithm. All GO enrichment results across stages are in **Supplemental Tables S2** and **S3**.

### Growth inhibition assay on K13^C580^ and K13^580Y^

K13^C580^ and K13^580Y^ cell lines were synchronized to the ring stage by treatment with 5% sorbitol for at least two successive replication cycles and cultured to the ring stage. Parasites were diluted with O+ RBCs to 0.4% parasitemia and 2% hematocrit in incomplete media (RPMI 1640 medium containing 25 mM HEPES and 2.0g/L sodium bicarbonate) supplemented with 0.5% AlbuMax II, 367.35µM hypoxanthine, and 25µg/mL gentamicin. Polyclonal mouse anti-GARP antiserum (10% v/v) (44) or normal mouse sera (10% v/v) was added to the cultures in 96-well round-bottom microtiter plates. After a 72-hour incubation, cultures were stained with Hoechst nucleic acid stain to quantify parasitemia using a CytoFLEX flow cytometer.

## RESULTS

### Experimental Design and Baseline Differential Expression (DE) between K13^580Y^ and K13^C580^

To study the transcriptomic profiles of *P. falciparum* cells in response to DHA exposure, we isolated coisogenic non-synchronized K13^C580^ and K13^580Y^ parasites after 48hrs in culture, and profiled the single cell transcriptome at different timepoints after DHA treatment (**Figure 1A**). The genetic difference between K13^580Y^ and K13^C580^ was verified using molecular inversion probes targeting multiple drug resistant markers (**Supplemental Methods** and **Supplemental Table S4**).

We first examined baseline DE prior to treatment (Time 0h, **Figure 1A**). Ring staged K13^580Y^ compared to K13^C580^ showed upregulation of 400 genes and downregulation of 219 genes. GO analysis detected enrichment of pathways related to parasite-erythrocyte interaction among upregulated genes (**Figure 1B** and **1C**), including erythrocyte aggregation, microvascular adhesion and receptor binding, represented by *GARP* (*45*), *RIFINS* (*PF3D7_0300500/PF3D7_1479400*), *FIKK5* (*46*)*, LyMP* (47)*, MESA* (*48*) and *Pf332* (*49*). Glycolytic pathways (*fructose-bisphosphate aldolase* (*FBPA*)) and food vacuole metabolism (*ornithine aminotransferase* (*OAT*) (50), *HSP60* and *HYP12*) were enriched in upregulated genes while mitochondrial ribosomal RNA genes (*LSUD* and *LSUF*) and apicoplast genes (*ClpM*) were downregulated, compared to K13^C580^ parasites. *K13* and *Kelch13 interacting candidate 2* (*KIC2*) (25) were significantly downregulated while *KIC9* was significantly upregulated in ring staged K13^580Y^ compared to K13^C580^ (**Supplemental Table S1**).

DE analysis between trophozoite staged K13^580Y^ vs. K13^C580^ (**Figure 1D** and **1E**) had greater DE differences relative to rings, with 201 and 1119 significantly up and down regulated genes, respectively. Similar to ring staged parasites, parasite-erythrocyte interaction pathways were enriched in upregulated genes. Variant surface antigen gene families were highly upregulated. Metabolic pathways were downregulated, including those in the mitochondria (*COX2A*, *COX17* and *COA5*) and apicoplast, such as isoprenoid pathway member *1-cys peroxiredoxin* (*AOP*) (51) and dolichol pathway component *polyprenol reductase* (*PPRD*) (52). *KIC3* was significantly downregulated in trophozoite staged K13^580Y^ compared to K13^C580^ (**Supplemental Table S1**).

### Differential Expression comparing DHA exposure in K13^C580^ and K13^580Y^

To quantify DHA perturbation across the malaria stages, the MELD algorithm compares DHA treatment to control over timepoints (see **Supplemental Methods**) (53). MELD score per cell, named the DHA likelihood, ranges from 0 to 1, i.e. from less to more perturbed by DHA treatment. Visualization of the distribution of DHA likelihoods across the lifecycle using pseudotime, calculated by slingshot (41), showed similar responses to DHA treatment in both strains, where there are separate transcriptional states based on treatment condition across the lifecycle (**Supplemental Figure S14**). As expected, DHA treated cells had a higher DHA likelihood than DMSO treated cells.

When focusing on stage specific DE, we isolated cell populations with the highest DHA likelihood within each timepoint using vertex frequency clustering (VFC) (53) (**Supplemental Figures S18-S22**). We separated these highly enriched clusters of high DHA likelihood cells (enriched in DHA treated cells) to compare with clusters of cells with low DHA likelihood (enriched in DMSO treated cells) to better examine drug effects in rings and trophozoites (**Supplemental Figures S18** and **S19**). VFC allowed for the isolation of cells prototypical for the effects of the DHA or DMSO treatment conditions. Direct DE results without MELD were comparable per stage (**Supplemental Figures S22-S24**).

Comparing DHA vs. DMSO treatment K13^C580^ rings across timepoints (**Figure 2A** and **2B**), the number of differentially upregulated genes increased as treatment continued across the timepoints (2h: 116 genes, 4h: 189 genes, 6h: 331 genes). GO analysis utilizing the significantly differentially expressed genes showed consistent downregulation over time of glycolytic genes (*ENO* and *LDH*), cell-cell adhesion (*var* genes *PF3D7_0200100*/*PF3D7_0712400*), food vacuole metabolism (*HRPIII*) and host cell surface pathways (*GARP*) (**Figure 2B**). Upregulated genes were enriched in ribosomal processing pathways (U3 ribonucleoprotein *IMP3*, *NOL10*) at 6h and nuclear genes (*Topoisomerase I*, histone H3, histone H2A) at 2h and 4h. There was sustained upregulation across timepoints in genes that assist in EMP1 loading (*PTP3*, *PTP4*, *PTP7*) and are exported proteins in the host cell membrane (*EVP1* (*54*), *PfG174* (*55*)*, exported lipase 2* (*XL2*) (56), *PfD80* (*57*) and *epoxide hydrolase 2* (*EH2*) (58)) (**Figure 2B**). Fatty acid metabolic gene *ACS10* was significantly upregulated across timepoints in K13^C580^ rings, while *K13* and *KIC5* were significantly upregulated at 4h and 6h with *KIC1* upregulated only at 6h (**Supplemental Table S1**).

**Figure 2:**
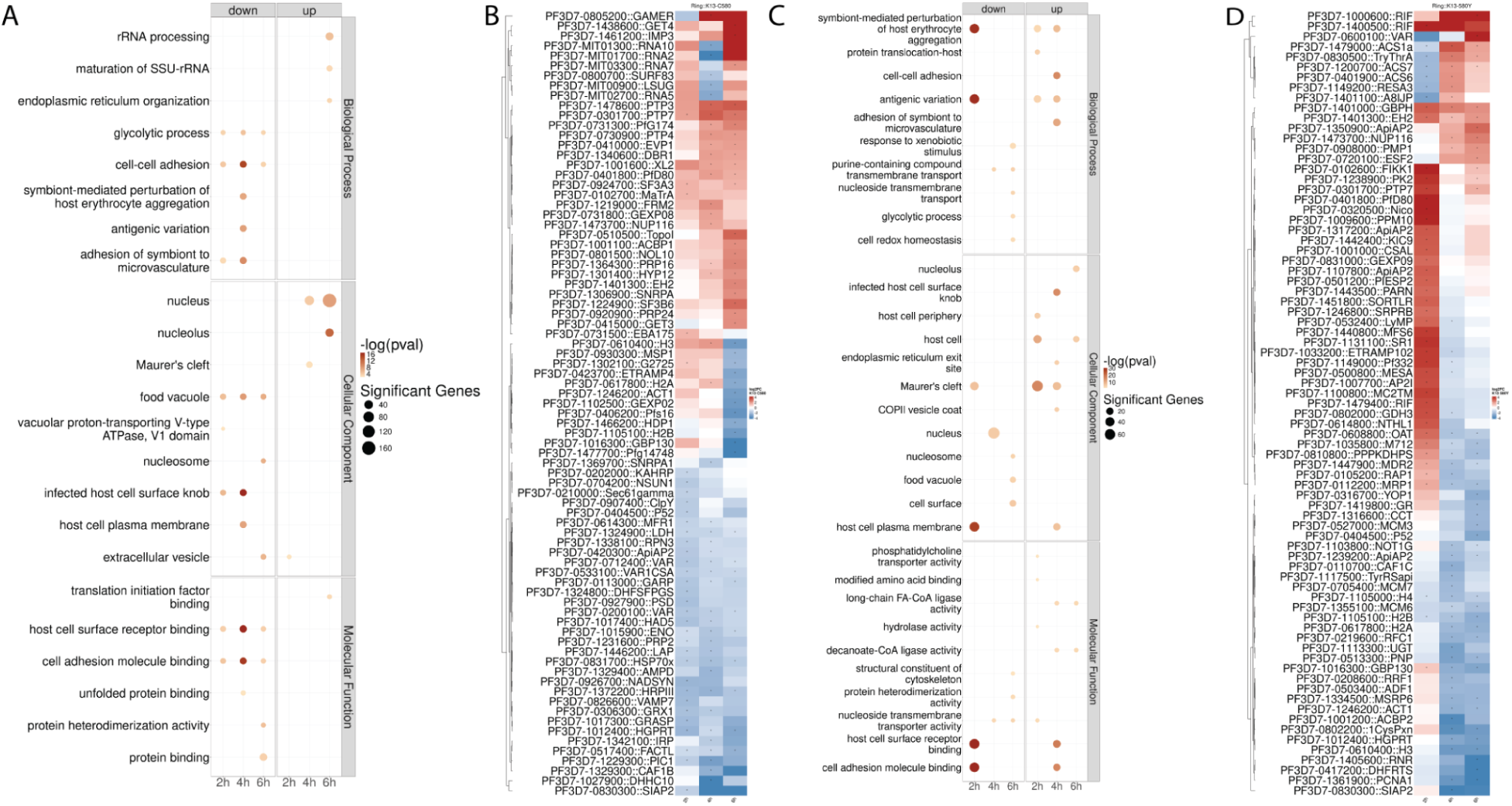
DHA Induced Upregulation of Exported Protein Genes, Downregulation of Glycolysis and Later Upregulation of Fatty Acid Metabolism in K13^C580^ and K13^580Y^ Rings. GO analysis was performed over the entire dataset in all differentially expressed genes in the selected VFC subcluster comparisons of K13^C580^ **(A)** or K13^580Y^ **(C)** trophozoites, with the number of significant genes that support that GO Term being the size of the circles (topGO p value < 0.01) and the color being the magnitude of the negative log of the topGO p-value. GO terms were filtered to the top 5 significant terms per timepoint. The union of the top 30 differentially expressed genes (by absolute log2FC) per time point between the VFC subclusters in the highest and lowest DHA likelihoods in K13^C580^ **(B)** or K13^580Y^ **(D)** trophozoites. The log2FC per timepoint is plotted. The top 5 hypervariable genes by absolute log2FC were selected for visualization for each heatmap. An asterisk indicates genes that were significantly differentially expressed at an absolute log2FC > 1 and a BH-adjusted p-value < 0.001. Red boxes are upregulated genes, white boxes indicate genes that were not differentially expressed, and blue boxes are downregulated genes.

Whereas in DHA vs. DMSO treated K13^580Y^ rings across timepoints (**Figure 2C** and **2D**), the greatest number of DE genes was at 2h, which then decreased over time (2h: 454 genes, 4h: 255 genes, 6h: 221 genes). At 2h, there was an initial upregulation of erythrocyte cell surface genes (*LyMP*, *Pf332, PfD80, MESA, FIKK1, ETRAMP102, PIESP2* (59) and *GEXP09*). At 4h and 6h post treatment, there was an upregulation of acyl-CoA synthetase genes involved in fatty acid utilization (*ACS1a*, *ACS6*, *ACS7* and *ACS10* (at 6h only, **Supplemental Table S1**)), and downregulation of nuclear pathways (*H2A*, *H2B*, *H3*, *H4* and *CAF1C*). *K13* and *KICs* were significantly upregulated; *KIC9* at 2h, *KIC4* at 4h, and *K13, KIC5* and *KIC6* at 4h and 6h (**Supplemental Table S1**).

### Fine Grained Differential Expression in Trophozoite Staged K13^C580^ or K13^580Y^

Comparing DHA or DMSO treatment in K13^C580^ trophozoites over time (**Figure 3A** and **3B**), the number of DE genes increased from 2h (77 genes) to similar levels at 4h (109 genes), and 6h (106 genes). At the 2 and 4 hour timepoints, top GO terms in each category were enriched in downregulated pathways consistent with slower metabolism: translation pathways (*elongation factor 2* (*eEF2*), *elongation factor 1-gamma* (*EF1gamma*) and *RPL39*), glycolysis (*ENO* and *triosephosphate isomerase* (*TIM*)), purine metabolism (*adenosine deaminase* (*ADA*), *nucleoside transporter 1 (NT1*), *nucleoside transporter 4* (*NT4*) and *PNP*) and fatty acid metabolism (*ACBP2* at 2h). At 2h and 4h post DHA treatment, artesunate inducible gene *PArt* (*60,61*) and genes involved with protein export (*FIKK12, PfD80, EMP3, PTP4/PTP7, FIRA*) were upregulated. At 6h post DHA treatment, there was continued upregulation of protein export including genes such as *RIFIN PF3D7_0115200*, *SPATR* (*62*)*, RAP1, AMA1* (*63*), *exported protein family 1* (*EPF1*) (64) and *C3H1 zinc finger 1* (*CZIF1*) (65). *KIC1* and *KIC9* were significantly upregulated at 2h and 4h, while *KIC10* was significantly upregulated at 6h (**Supplemental Table S1**).

**Figure 3:**
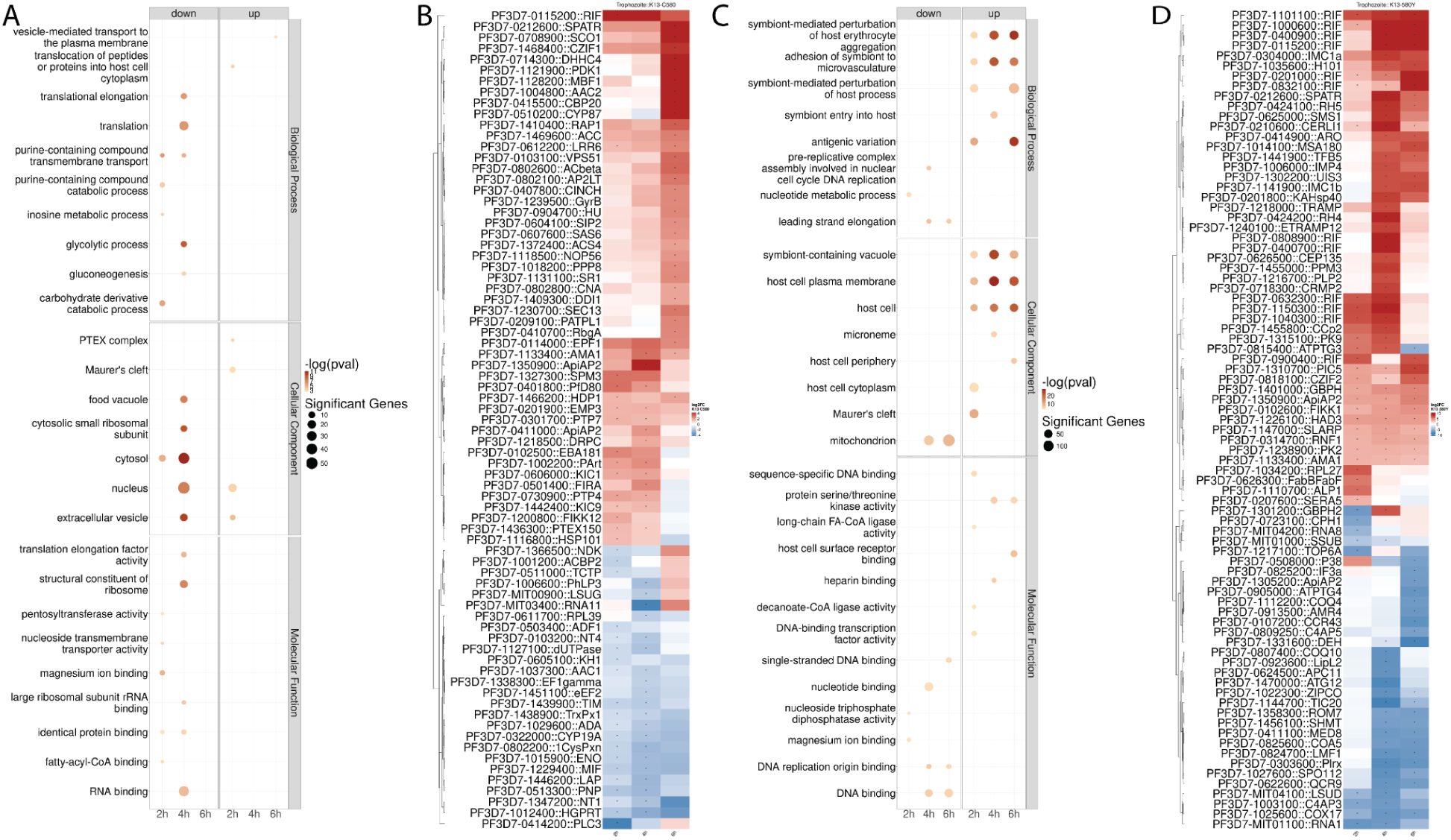
DHA Treatment Induced Protein Export Genes in K13^C580^ and K13^580Y^ Trophozoites, with Broad Downregulation of Mitochondrial and Apicoplast Pathways in K13^580Y^ Trophozoites. GO analysis was performed over all differentially expressed genes in the selected VFC subcluster comparisons of K13^C580^ **(A)** or K13^580Y^ **(C)** trophozoites, with the number of significant genes that support that GO Term being the size of the circles (topGO p value < 0.01) and the color being the magnitude of the negative log of the topGO p-value. GO terms were filtered to the top 5 significant terms per timepoint. The union of the top 30 differentially expressed genes (by absolute log2FC) per time point between the VFC subclusters in the highest and lowest DHA likelihoods in K13^C580^ **(B)** or K13^580Y^ **(D)** trophozoites. The top 5 hypervariable genes by absolute log2FC were selected for visualization for each heatmap. An asterisk indicates genes that were significantly differentially expressed at an absolute log2FC > 1 and a BH-adjusted p-value < 0.001. Red boxes are upregulated genes, white boxes indicate genes that were not differentially expressed, and blue boxes are downregulated genes.

After DHA treatment in K13^580Y^ trophozoites over time (**Figure 3C** and **3D**), there was greater induction of transcription compared to the DHA treated K13^C580^ trophozoites (2h: 569 genes, 4h: 1074 genes, 6h: 1244 DE genes). Across time, there was a strong and broad upregulation of genes related to parasite-erythrocyte interaction, invasion and adhesion to microvasculature, including *RIFINs PF3D7_0900400*/*PF3D7_1040300, SPATR, FIKK1, AMA1, CZIF2, merozoite surface protein* (*H101*), *cytosolically exposed rhoptry leaflet interacting protein 2* (*CERLI*)(66)*, reticulocyte-binding protein homologue 4 (RH4)/RH5* (67)*, thrombospondin-related apical merozoite protein (TRAMP)* (*68*), *ETRAMP12,* and heat shock protein *KAHsp40* (*69*).

Mitochondrial and apicoplast pathways were downregulated, containing genes such as *C4AP3/C4AP5, COQ4/COQ10, COX17, cytochrome c oxidase assembly factor 5* (*COA5*)*, QCR9, LSUD*, *lipoate-protein ligase 2* (*LIPL2)* (*70*) and *AMR4* (*71*). Additionally, there was downregulation of genes related to translation at 4h and 6h, such as *EF1gamma* and *eEF2*. *K13, KIC1, KIC5, KIC6, KIC7, KIC4, KIC9* was significantly upregulated across timepoints, *KIC2* was significantly upregulated at 4h and 6h, and *KIC10* was significantly downregulated at 6h (**Supplemental Table S1**).

### Anti-GARP kills K13^C580^ *and* K13^580Y^ *in vitro*

Given GARP expression was particularly elevated in untreated K13^580Y^ compared to K13^C580^ rings (**Figure 1B**), we evaluated if this elevated expression was a result of a functional or aberrant accumulation of the protein in the membrane. We performed growth-inhibition assays (GIAs) using polyclonal mouse anti-GARP antiserum (44). We found statistically significant growth inhibition of both K13^C580^ or K13^580Y^ lines after exposure to anti-GARP (**Figure 4**), consistent with functionality of the pathway in both lines and the upregulation of GARP in the K13^580Y^ line (**Figure 4**). This result suggests that other elevated surface proteins, like GARP, could remain functionally intact and may contribute to the DHA response.

**Figure 4:**
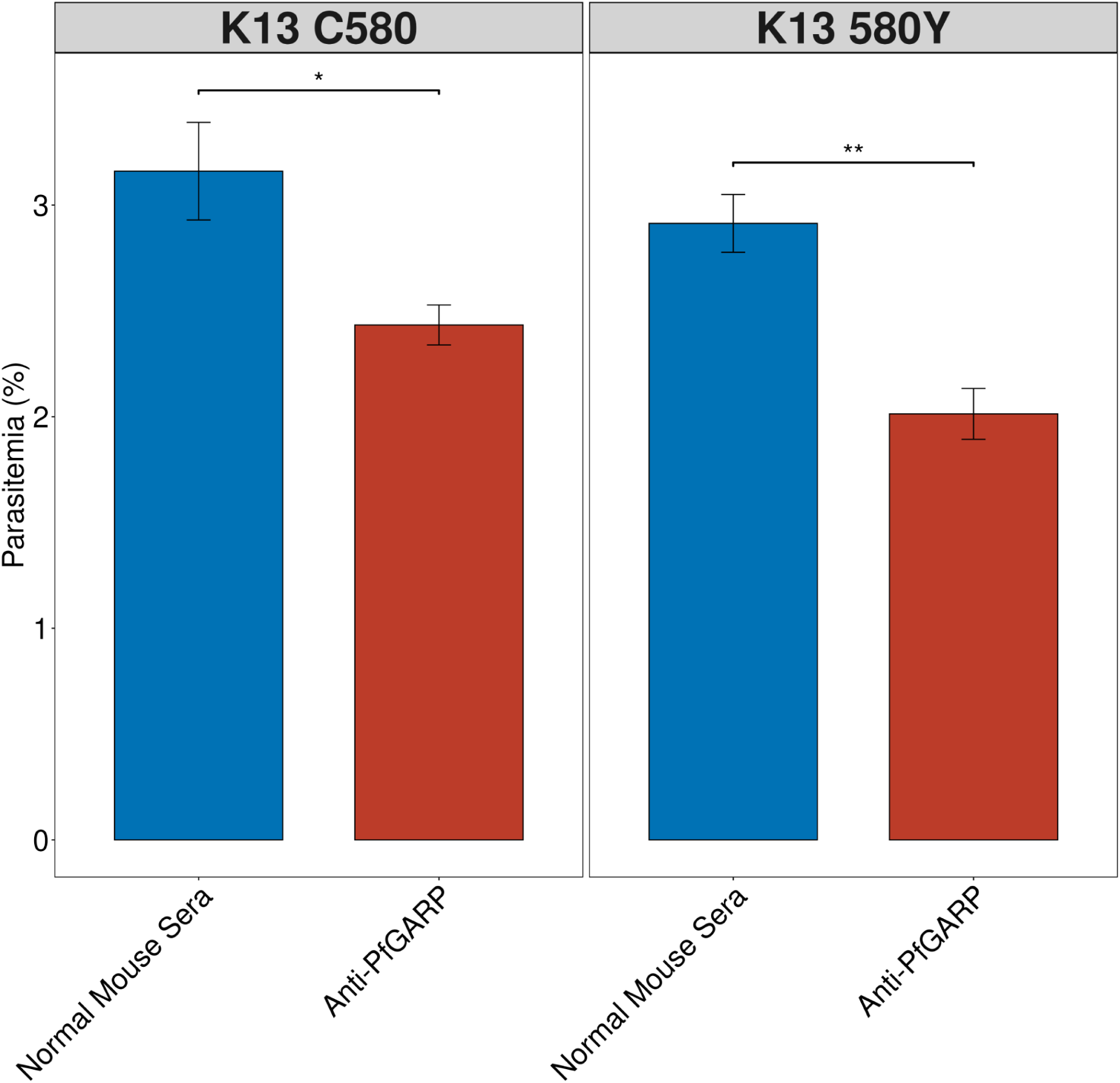
Anti-PfGARP inhibits parasite growth in both artemisinin sensitive and resistant parasites. Artemisinin sensitive (K13^C580^) and resistant (K13^580Y^) parasites were synchronized to the ring stage, adjusted to 0.4% parasitemia and 2% hematocrit and incubated with anti-PfGARP (10% v/v) or normal mouse sera (10% v/v). Parasites were cultured for 72 hours followed by quantification of parasitemia by flow cytometry. Bars represent the mean of three technical replicates and error bars represent mean +/- standard deviation. Results representative of 3 independent experiments (unpaired t-test with bonferroni adjustment, *p<0.5, **p<0.01).

## DISCUSSION

To date, studies of *Plasmodium falciparum* transcriptional responses to artemisinin have largely relied on bulk RNA sequencing of synchronized parasites, limiting resolution across developmental stages and obscuring cell-to-cell heterogeneity (16,72–75). Here, we leveraged single-cell RNA sequencing to comprehensively profile the transcriptional landscape of artemisinin-sensitive (K13^C580^) and artemisinin-resistant (K13^580Y^) coisogenic parasites following dihydroartemisinin (DHA) exposure. This approach revealed stage-specific and strain-specific responses. In comparing K13^580Y^ vs. K13^C580^ untreated rings, there was an upregulation of exported protein related genes (notably *GARP, MESA, Pf332*) and genes in metabolic pathways, with upregulated genes enriched in host cell modification and virulence functions (**Figure 1**). Differential expression in trophozoites showed downregulation of mitochondrial and apicoplast genes when comparing K13^580Y^ vs. K13^C580^. This expression pattern could indicate that K13^580Y^ parasites have adapted a transcriptional program to increase exported protein pathways, while reducing mitochondrial sources of reactive oxygen species, to survive drug treatment at the tradeoff of fitness (75,76).

Using MELD, we quantified the extent of experimental perturbation across the lifecycle. We separated out populations of rings and trophozoites that were most and least perturbed by DHA treatment to perform DE analysis. In K13^C580^ and K13^580Y^ ring stages, there was upregulation of exported protein pathways and genes involved in fatty acid utilization (**Figure 2A** and **Supplemental Table S1**). Fatty-acid metabolism is hypothesized to be an alternative energy source to sustain parasite viability upon DHA treatment (77,78). Pires *et al.* proposed a stress response upon DHA treatment that consists of the “exportome” of secreted parasite proteins (79) acting in synergy to facilitate lipid transport for fatty acid metabolism (77). Both lines showed downregulation of glycolytic processes and upregulation of exported protein and fatty acid metabolic genes (**Figure 2)**. DHA treated ring staged parasites’ broad activation of exported proteins and increase in fatty acid metabolic pathway genes suggests a shift to fatty acid metabolism to respond to DHA induced stress. Future studies aiming to inhibit acyl-CoA synthetases, such as ACS10, may be a promising avenue to bolstering DHA treatment (80).

In both K13^C580^ and K13^580Y^ DHA-treated trophozoites, there was an upregulation of exported protein related genes and invasion pathway genes. Additionally, there was a downregulation of metabolism and translation in both strains (**Figure 3**). These results support previous reports of translational down-regulation after DHA treatment (73). Interestingly, both DHA-treated K13^C580^ and K13^580Y^ trophozoites showed downregulation of genes involved in glycolysis, purine metabolism and fatty acid metabolism (**Figure 3** and **Supplemental Table S1**). Whereas only in DHA-treated K13^580Y^ trophozoites, we saw broad downregulation of mitochondrial and apicoplast pathway gene expression. This result suggests that the K13 mutation may alter different metabolic pathways in response to DHA treatment, similar to prior work showing differential mitochondrial pathway use in sensitive vs. resistant lines (73).

Throughout our analysis, we saw differential upregulation or downregulation of transcriptional regulators, such as *ApiAP2* family genes (**Figures 2** and **3**). Recently, Yu *et al.* identified epigenetic modulators that contribute to ring stage survival after DHA treatment, notably the histone acetyltransferase *MYST* (81). Our findings showing altered expression of transcription factor activity underscores the importance to further investigate transcriptional regulation of artemisinin response.

The K13 protein exhibits sequence similarity to a class of Kelch/BTB/POZ ubiquitination adaptors and functions within the endocytosis pathway required for hemoglobin uptake from the host cell (19,82), which is critical for the activation of the endoperoxide bridge of ART (83). *K13* mutations in the propeller domain are thought to partially reduce the normal function of the K13 protein, which in turn is thought to diminish hemoglobin endocytosis and ART activation (25,26). Reduced hemoglobin uptake limits the availability of Fe^2+^ heme to activate ART and kill ring stage parasites via alkylation of proximal proteins, leading to reduced ART killing (83,84). At baseline, *K13* was downregulated in K13^580Y^ ring staged parasites compared to K13^C580^ K13’s function in facilitating hemoglobin entry has been shown to be aided by a complex of other proteins, known as the Kelch13 interacting candidates (KICs) (25). Downregulation of hemoglobin peptides upon DHA treatment occurs regardless of *K13* genotype (73). There was a broad upregulation of *KIC* genes that are localized to the K13 endocytosis compartment (such as *KIC4, KIC5* and *KIC7*) (85) across rings and trophozoites of both lines after DHA treatment, possibly as a compensatory measure to preserve hemoglobin endocytosis.

Differential expression in ring staged K13^580Y^ vs. K13^C580^ showed upregulated levels of GARP in K13^580Y^ (**Figure 1C**). GARP was recently identified as a therapeutic target (44), while its function remains to be characterized. We observed elevated expression of GARP in K13^580Y^ compared to K13^C580^ before treatment, possibly as a result of compensatory increase in the expression of the “exportome” (86). Given gene expression changes can correlate with function or represent dysregulation (e.g. increased turnover), we examined whether GARP still retained function as a therapeutic target, finding statistically significant anti-GARP killing of both K13^C580^ and K13^580Y^ parasites (**Figure 4**). This suggests that targeting GARP and likely other surface proteins may be effective therapy for K13 mutant parasites.

Our study provides a view of the single-cell transcriptional landscape underlying artemisinin response in *Plasmodium falciparum*, revealing both shared stress pathways and resistance-associated adaptations. Together, these findings not only advance our understanding of the molecular basis of artemisinin resistance but also provide a framework for rational design of next-generation therapies that exploit vulnerabilities in resistant parasites. By leveraging single-cell RNA sequencing, we demonstrate that DHA treated parasites deploy a distinct transcriptional program characterized by enhanced protein export and metabolic flexibility, including fatty acid utilization, while downregulating organellar functions. Importantly, the inhibition of GARP and its possible role in sustaining parasite growth through lipid metabolism underscores the potential of targeting exportome components to overcome resistance. Future work integrating transcriptomic, proteomic, and functional studies will be critical to better assess and translate these insights into effective interventions against evolving malaria resistance.

## Data availability

All sequence data has been submitted to SRA (PRJNA1049964). Code is available at https://github.com/sconnelly007/K13_mt_v_wt_sc.

## Author contributions

SVC, CIO, CC, JAB, JK, and JJJ designed the research study. CIO, CC, IP, DS, NR, and JZ conducted the experiments. SVC and CIO analyzed the data. SVC, CIO, JAB and JJJ wrote the manuscript. All authors reviewed and revised the manuscript.

## Ethics Statement

This research does not constitute human subjects research.

## Generative AI Statement

ChatGPT was consulted for the editing for select paragraphs and for editing of code used for analysis.

## References

1. World Health Organization. World malaria report 2025 [Internet]. World Health Organization; 2025 Dec [cited 2026 Jan 1]. Available from: https://www.who.int/publications/i/item/9789240117822

2. Teuscher F, Gatton ML, Chen N, Peters J, Kyle DE, Cheng Q. Artemisinin-induced dormancy in Plasmodium falciparum: Duration, recovery rates, and implications in treatment failure. J Infect Dis. 2010 Nov;202(9):1362–8.

3. Noedl H, Se Y, Schaecher K, Smith BL, Socheat D, Fukuda MM, et al. Evidence of artemisinin-resistant malaria in western Cambodia. N Engl J Med. 2008 Dec 11;359(24):2619–20.

4. Imwong M, Dhorda M, Myo Tun K, Thu AM, Phyo AP, Proux S, et al. Molecular epidemiology of resistance to antimalarial drugs in the Greater Mekong subregion: an observational study. Lancet Infect Dis. 2020 Dec;20(12):1470–80.

5. Tacoli C, Gai PP, Bayingana C, Sifft K, Geus D, Ndoli J, et al. Artemisinin resistance-associated K13 polymorphisms of Plasmodium falciparum in southern Rwanda, 2010-2015. Am J Trop Med Hyg. 2016 Nov 2;95(5):1090–3.

6. Ashley EA, Dhorda M, Fairhurst RM, Amaratunga C, Lim P, Suon S, et al. Spread of artemisinin resistance in Plasmodium falciparum malaria. N Engl J Med. 2014 Jul 31;371(5):411–23.

7. Fola AA, Feleke SM, Mohammed H, Brhane BG, Hennelly CM, Assefa A, et al. Plasmodium falciparum resistant to artemisinin and diagnostics have emerged in Ethiopia. Nat Microbiol. 2023 Aug 28;8(10):1911–9.

8. Juliano JJ, Giesbrecht DJ, Simkin A, Fola AA, Lyimo BM, Pereus D, et al. Prevalence of mutations associated with artemisinin partial resistance and sulfadoxine-pyrimethamine resistance in 13 regions in Tanzania in 2021: a cross-sectional survey. Lancet Microbe. 2024 Oct;5(10):100920.

9. Balikagala B, Fukuda N, Ikeda M, Katuro OT, Tachibana SI, Yamauchi M, et al. Evidence of artemisinin-resistant malaria in Africa. N Engl J Med. 2021 Sep 23;385(13):1163–71.

10. Conrad MD, Asua V, Garg S, Giesbrecht D, Niaré K, Smith S, et al. Evolution of Partial Resistance to Artemisinins in Malaria Parasites in Uganda. N Engl J Med. 2023 Aug 24;389(8):722–32.

11. Moriarty LF, Nkoli PM, Likwela JL, Mulopo PM, Sompwe EM, Rika JM, et al. Therapeutic Efficacy of Artemisinin-Based Combination Therapies in Democratic Republic of the Congo and Investigation of Molecular Markers of Antimalarial Resistance. Am J Trop Med Hyg. 2021 Sep 7;105(4):1067–75.

12. Dimbu PR, Horth R, Cândido ALM, Ferreira CM, Caquece F, Garcia LEA, et al. Continued Low Efficacy of Artemether-Lumefantrine in Angola in 2019. Antimicrob Agents Chemother [Internet]. 2021 Jan 20;65(2). Available from: 10.1128/AAC.01949-20

13. Ebong C, Sserwanga A, Namuganga JF, Kapisi J, Mpimbaza A, Gonahasa S, et al. Efficacy and safety of artemether-lumefantrine and dihydroartemisinin-piperaquine for the treatment of uncomplicated Plasmodium falciparum malaria and prevalence of molecular markers associated with artemisinin and partner drug resistance in Uganda. Malar J. 2021 Dec 24;20(1):484.

14. Gansané A, Moriarty LF, Ménard D, Yerbanga I, Ouedraogo E, Sondo P, et al. Anti-malarial efficacy and resistance monitoring of artemether-lumefantrine and dihydroartemisinin-piperaquine shows inadequate efficacy in children in Burkina Faso, 2017-2018. Malar J. 2021 Jan 19;20(1):48.

15. Connelly SV, Muller JG, Ali M, Ngasala BE, Hassan W, Mohamed B, et al. Artemisinin partial resistance mutations in Zanzibar and Tanzania suggest regional spread and African origins, 2023. J Infect Dis [Internet]. 2025 Aug 13;(jiaf431). Available from: 10.1093/infdis/jiaf431/8233845

16. Dondorp AM, Nosten F, Yi P, Das D, Phyo AP, Tarning J, et al. Artemisinin resistance in Plasmodium falciparum malaria. N Engl J Med. 2009 Jul 30;361(5):455–67.

17. Björkman A, Gil P, Alifrangis M. Alarming Plasmodium falciparum resistance to artemisinin-based combination therapy in Africa: the critical role of the partner drug. Lancet Infect Dis. 2024 Sep;24(9):e540–1.

18. Dhorda M, Kaneko A, Komatsu R, Kc A, Mshamu S, Gesase S, et al. Artemisinin-resistant malaria in Africa demands urgent action. Science. 2024 Jul 19;385(6706):252–4.

19. Ariey F, Witkowski B, Amaratunga C, Beghain J, Langlois AC, Khim N, et al. A molecular marker of artemisinin-resistant Plasmodium falciparum malaria. Nature. 2014 Jan 2;505(7481):50–5.

20. Straimer J, Gnädig NF, Witkowski B, Amaratunga C, Duru V, Ramadani AP, et al. Drug resistance. K13-propeller mutations confer artemisinin resistance in Plasmodium falciparum clinical isolates. Science. 2015 Jan 23;347(6220):428–31.

21. MalariaGEN Plasmodium falciparum Community Project. Genomic epidemiology of artemisinin resistant malaria. Elife. 2016 Mar 4;5:e08714.

22. Ménard D, Khim N, Beghain J, Adegnika AA, Shafiul-Alam M, Amodu O, et al. A Worldwide Map of Plasmodium falciparum K13-Propeller Polymorphisms. N Engl J Med. 2016 Jun 23;374(25):2453–64.

23. World Health Organization. Compendium of molecular markers for antimalarial drug resistance [Internet]. 2025. Available from: https://www.who.int/publications/m/item/compendium-of-molecular-markers-for-antimalarial-drug-resistance

24. Ishengoma DS, Gosling R, Martinez-Vega R, Beshir KB, Bailey JA, Chimumbwa J, et al. Urgent action is needed to confront artemisinin partial resistance in African malaria parasites. Nat Med. 2024 Jul;30(7):1807–8.

25. Birnbaum J, Scharf S, Schmidt S, Jonscher E, Hoeijmakers WAM, Flemming S, et al. A Kelch13-defined endocytosis pathway mediates artemisinin resistance in malaria parasites. Science. 2020 Jan 3;367(6473):51–9.

26. Yang T, Yeoh LM, Tutor MV, Dixon MW, McMillan PJ, Xie SC, et al. Decreased K13 abundance reduces hemoglobin catabolism and proteotoxic stress, underpinning artemisinin resistance. Cell Rep. 2019 Nov 26;29(9):2917–28.e5.

27. Stokes BH, Dhingra SK, Rubiano K, Mok S, Straimer J, Gnädig NF, et al. Plasmodium falciparum K13 mutations in Africa and Asia impact artemisinin resistance and parasite fitness. Elife. 2021 Jul 19;10(e66277):e66277.

28. Klonis N, Xie SC, McCaw JM, Crespo-Ortiz MP, Zaloumis SG, Simpson JA, et al. Altered temporal response of malaria parasites determines differential sensitivity to artemisinin. Proc Natl Acad Sci U S A. 2013 Mar 26;110(13):5157–62.

29. Mok S, Imwong M, Mackinnon MJ, Sim J, Ramadoss R, Yi P, et al. Artemisinin resistance in Plasmodium falciparum is associated with an altered temporal pattern of transcription. BMC Genomics. 2011 Aug 3;12(1):391.

30. Howick VM, Russell AJC, Andrews T, Heaton H, Reid AJ, Natarajan K, et al. The Malaria Cell Atlas: Single parasite transcriptomes across the complete Plasmodium life cycle. Science [Internet]. 2019 Aug 23;365(6455). Available from: 10.1126/science.aaw2619

31. Dogga SK, Rop JC, Cudini J, Farr E, Dara A, Ouologuem D, et al. A single cell atlas of sexual development in Plasmodium falciparum. Science. 2024 May 3;384(6695):eadj4088.

32. Nötzel C, Kafsack BFC. PfD123 modulates K13-mediated survival and recovery after artemisinin exposure [Internet]. bioRxiv. 2022 [cited 2023 Mar 3]. p. 2022.01.27.476788. Available from: https://www.biorxiv.org/content/10.1101/2022.01.27.476788

33. Gierahn TM, Wadsworth MH 2nd, Hughes TK, Bryson BD, Butler A, Satija R, et al. Seq-Well: portable, low-cost RNA sequencing of single cells at high throughput. Nat Methods. 2017 Apr;14(4):395–8.

34. Hughes TK, Wadsworth MH 2nd, Gierahn TM, Do T, Weiss D, Andrade PR, et al. Second-Strand Synthesis-based massively parallel scRNA-seq reveals cellular states and molecular features of human inflammatory skin pathologies. Immunity. 2020 Oct 13;53(4):878–94.e7.

35. Aicher TP, Carroll S, Raddi G, Gierahn T, Wadsworth MH 2nd, Hughes TK, et al. Seq-Well: A sample-efficient, portable picowell platform for massively parallel single-cell RNA sequencing. Methods Mol Biol. 2019;1979:111–32.

36. Schuster FL. Cultivation of plasmodium spp. Clin Microbiol Rev. 2002 Jul;15(3):355–64.

37. Brown AC, Moore CC, Guler JL. Cholesterol-dependent enrichment of understudied erythrocytic stages of human Plasmodium parasites. Sci Rep. 2020 Mar 12;10(1):4591.

38. Macosko EZ, Basu A, Satija R, Nemesh J, Shekhar K, Goldman M, et al. Highly Parallel Genome-wide Expression Profiling of Individual Cells Using Nanoliter Droplets. Cell. 2015 May 21;161(5):1202–14.

39. Butler A, Hoffman P, Smibert P, Papalexi E, Satija R. Integrating single-cell transcriptomic data across different conditions, technologies, and species. Nat Biotechnol. 2018 May;36(5):411–20.

40. Neavin D, Senabouth A, Arora H, Lee JTH, Ripoll-Cladellas A, sc-eQTLGen Consortium, et al. Demuxafy: improvement in droplet assignment by integrating multiple single-cell demultiplexing and doublet detection methods. Genome Biol. 2024 Apr 15;25(1):94.

41. Street K, Risso D, Fletcher RB, Das D, Ngai J, Yosef N, et al. Slingshot: cell lineage and pseudotime inference for single-cell transcriptomics. BMC Genomics [Internet]. 2018 Dec;19(1). Available from: 10.1186/s12864-018-4772-0

42. Alexa A, Rahnenfuhrer J. topGO: enrichment analysis for gene ontology. R package version [Internet]. 2010; Available from: https://scholar.google.com/citations?user=IIVQsKIAAAAJ&hl=en&oi=sra

43. Simmons C, Gibbons J, Wang C, Pires CV, Zhang M, Siddiqui F, et al. A novel Modulator of Ring Stage Translation (MRST) gene alters artemisinin sensitivity in Plasmodium falciparum. mSphere. 2023 May 23;8(4):e0015223.

44. Raj DK, Das Mohapatra A, Jnawali A, Zuromski J, Jha A, Cham-Kpu G, et al. Anti-PfGARP activates programmed cell death of parasites and reduces severe malaria. Nature. 2020 Jun;582(7810):104–8.

45. Almukadi H, Schwake C, Kaiser MM, Mayer DCG, Schiemer J, Baldwin MR, et al. Human erythrocyte band 3 is a host receptor for Plasmodium falciparum glutamic acid-rich protein. Blood. 2019 Jan 31;133(5):470–80.

46. Ward P, Equinet L, Packer J, Doerig C. Protein kinases of the human malaria parasite Plasmodium falciparum: the kinome of a divergent eukaryote. BMC Genomics. 2004 Oct 12;5(1):79.

47. Proellocks NI, Herrmann S, Buckingham DW, Hanssen E, Hodges EK, Elsworth B, et al. A lysine-rich membrane-associated PHISTb protein involved in alteration of the cytoadhesive properties of Plasmodium falciparum-infected red blood cells. FASEB J. 2014 Jul;28(7):3103–13.

48. Coppel RL, Lustigman S, Murray L, Anders RF. MESA is a Plasmodium falciparum phosphoprotein associated with the erythrocyte membrane skeleton. Mol Biochem Parasitol. 1988 Dec;31(3):223–31.

49. Hodder AN, Maier AG, Rug M, Brown M, Hommel M, Pantic I, et al. Analysis of structure and function of the giant protein Pf332 in Plasmodium falciparum. Mol Microbiol. 2009 Jan;71(1):48–65.

50. Gafan C, Wilson J, Berger LC, Berger BJ. Characterization of the ornithine aminotransferase from Plasmodium falciparum. Mol Biochem Parasitol. 2001 Nov;118(1):1–10.

51. Swift RP, Rajaram K, Elahi R, Liu HB, Prigge ST. Roles of Ferredoxin-Dependent Proteins in the Apicoplast of Plasmodium falciparum Parasites. MBio. 2021 Feb 22;13(1):e0302321.

52. Zimbres FM, Valenciano AL, Merino EF, Florentin A, Holderman NR, He G, et al. Metabolomics profiling reveals new aspects of dolichol biosynthesis in Plasmodium falciparum. Sci Rep. 2020 Aug 6;10(1):13264.

53. Burkhardt DB, Stanley JS 3rd, Tong A, Perdigoto AL, Gigante SA, Herold KC, et al. Quantifying the effect of experimental perturbations at single-cell resolution. Nat Biotechnol. 2021 May;39(5):619–29.

54. Tamez PA, Bhattacharjee S, van Ooij C, Hiller NL, Llinás M, Balu B, et al. An erythrocyte vesicle protein exported by the malaria parasite promotes tubovesicular lipid import from the host cell surface. PLoS Pathog. 2008 Aug 8;4(8):e1000118.

55. Vincensini L, Richert S, Blisnick T, Van Dorsselaer A, Leize-Wagner E, Rabilloud T, et al. Proteomic analysis identifies novel proteins of the Maurer’s clefts, a secretory compartment delivering Plasmodium falciparum proteins to the surface of its host cell. Mol Cell Proteomics. 2005 Apr;4(4):582–93.

56. Liu J, Fike KR, Dapper C, Klemba M. Metabolism of host lysophosphatidylcholine in Plasmodium falciparum-infected erythrocytes. Proc Natl Acad Sci U S A. 2024 Feb 20;121(8):e2320262121.

57. Jonsdottir TK, Counihan NA, Modak JK, Kouskousis B, Sanders PR, Gabriela M, et al. Characterisation of complexes formed by parasite proteins exported into the host cell compartment of Plasmodium falciparum infected red blood cells. Cell Microbiol. 2021 Aug;23(8):e13332.

58. Spillman NJ, Dalmia VK, Goldberg DE. Exported Epoxide Hydrolases Modulate Erythrocyte Vasoactive Lipids during Plasmodium falciparum Infection. MBio [Internet]. 2016 Oct 18;7(5). Available from: 10.1128/mBio.01538-16

59. Florens L, Liu X, Wang Y, Yang S, Schwartz O, Peglar M, et al. Proteomics approach reveals novel proteins on the surface of malaria-infected erythrocytes. Mol Biochem Parasitol. 2004 May;135(1):1–11.

60. Natalang O, Bischoff E, Deplaine G, Proux C, Dillies MA, Sismeiro O, et al. Dynamic RNA profiling in Plasmodium falciparum synchronized blood stages exposed to lethal doses of artesunate. BMC Genomics. 2008 Aug 18;9(1):388.

61. Deplaine G, Lavazec C, Bischoff E, Natalang O, Perrot S, Guillotte-Blisnick M, et al. Artesunate tolerance in transgenic Plasmodium falciparum parasites overexpressing a tryptophan-rich protein. Antimicrob Agents Chemother. 2011 Jun;55(6):2576–84.

62. Gupta R, Mishra A, Choudhary HH, Narwal SK, Nayak B, Srivastava PN, et al. Secreted protein with altered thrombospondin repeat (SPATR) is essential for asexual blood stages but not required for hepatocyte invasion by the malaria parasite Plasmodium berghei. Mol Microbiol. 2020 Feb;113(2):478–91.

63. Treeck M, Zacherl S, Herrmann S, Cabrera A, Kono M, Struck NS, et al. Functional analysis of the leading malaria vaccine candidate AMA-1 reveals an essential role for the cytoplasmic domain in the invasion process. PLoS Pathog. 2009 Mar;5(3):e1000322.

64. Mbengue A, Audiger N, Vialla E, Dubremetz JF, Braun-Breton C. Novel Plasmodium falciparum Maurer’s clefts protein families implicated in the release of infectious merozoites. Mol Microbiol. 2013 Apr;88(2):425–42.

65. Balbin JM, Heinemann GK, Yeoh LM, Gilberger TW, Armstrong M, Duffy MF, et al. Characterisation of PfCZIF1 and PfCZIF2 in Plasmodium falciparum asexual stages. Int J Parasitol. 2023 Jan;53(1):27–41.

66. Liffner B, Frölich S, Heinemann GK, Liu B, Ralph SA, Dixon MWA, et al. PfCERLI1 is a conserved rhoptry associated protein essential for Plasmodium falciparum merozoite invasion of erythrocytes. Nat Commun. 2020 Mar 16;11(1):1411.

67. Baum J, Chen L, Healer J, Lopaticki S, Boyle M, Triglia T, et al. Reticulocyte-binding protein homologue 5 - an essential adhesin involved in invasion of human erythrocytes by Plasmodium falciparum. Int J Parasitol. 2009 Feb;39(3):371–80.

68. Scally SW, Triglia T, Evelyn C, Seager BA, Pasternak M, Lim PS, et al. PCRCR complex is essential for invasion of human erythrocytes by Plasmodium falciparum. Nat Microbiol. 2022 Dec;7(12):2039–53.

69. Acharya P, Chaubey S, Grover M, Tatu U. An exported heat shock protein 40 associates with pathogenesis-related knobs in Plasmodium falciparum infected erythrocytes. PLoS One. 2012 Sep 7;7(9):e44605.

70. Günther S, Wallace L, Patzewitz EM, McMillan PJ, Storm J, Wrenger C, et al. Apicoplast lipoic acid protein ligase B is not essential for Plasmodium falciparum. PLoS Pathog. 2007 Dec;3(12):e189.

71. Meister TR, Tang Y, Pulkoski-Gross MJ, Yeh E. CaaX-like protease of cyanobacterial origin is required for complex Plastid biogenesis in malaria parasites. MBio [Internet]. 2020 Oct 6;11(5). Available from: 10.1128/mBio.01492-20

72. Zhu L, Tripathi J, Rocamora FM, Miotto O, van der Pluijm R, Voss TS, et al. The origins of malaria artemisinin resistance defined by a genetic and transcriptomic background. Nat Commun. 2018 Dec 4;9(1):5158.

73. Mok S, Stokes BH, Gnädig NF, Ross LS, Yeo T, Amaratunga C, et al. Artemisinin-resistant K13 mutations rewire Plasmodium falciparum’s intra-erythrocytic metabolic program to enhance survival. Nat Commun. 2021 Jan 22;12(1):530.

74. Mok S, Ashley EA, Ferreira PE, Zhu L, Lin Z, Yeo T, et al. Drug resistance. Population transcriptomics of human malaria parasites reveals the mechanism of artemisinin resistance. Science. 2015 Jan 23;347(6220):431–5.

75. Zhu L, van der Pluijm RW, Kucharski M, Nayak S, Tripathi J, White NJ, et al. Artemisinin resistance in the malaria parasite, Plasmodium falciparum, originates from its initial transcriptional response. Commun Biol. 2022 Mar 28;5(1):274.

76. Behrens HM, Schmidt S, Henshall IG, López-Barona P, Peigney D, Sabitzki R, et al. Impact of different mutations on Kelch13 protein levels, ART resistance, and fitness cost in Plasmodium falciparum parasites. MBio. 2024 May 3;15(6):e0198123.

77. Pires CV, Oberstaller J, Wang C, Casandra D, Zhang M, Chawla J, et al. Chemogenomic profiling of a Plasmodium falciparum transposon mutant library reveals shared effects of dihydroartemisinin and bortezomib on lipid metabolism and exported proteins. Microbiol Spectr. 2023 Jun 15;11(3):e0501422.

78. Chen N, LaCrue AN, Teuscher F, Waters NC, Gatton ML, Kyle DE, et al. Fatty acid synthesis and pyruvate metabolism pathways remain active in dihydroartemisinin-induced dormant ring stages of Plasmodium falciparum. Antimicrob Agents Chemother. 2014 Aug;58(8):4773–81.

79. Hiller NL, Bhattacharjee S, van Ooij C, Liolios K, Harrison T, Lopez-Estraño C, et al. A host-targeting signal in virulence proteins reveals a secretome in malarial infection. Science. 2004 Dec 10;306(5703):1934–7.

80. Bopp S, Pasaje CFA, Summers RL, Magistrado-Coxen P, Schindler KA, Corpas-Lopez V, et al. Potent acyl-CoA synthetase 10 inhibitors kill Plasmodium falciparum by disrupting triglyceride formation. Nat Commun. 2023 Mar 16;14(1):1455.

81. Yu X, He J, Wang C, Mu J, Chen X, Zhao Y, et al. Epigenetically conferred ring-stage survival in Plasmodium falciparum against artemisinin treatment. Nat Commun. 2025 Aug 28;16(1):8037.

82. Xie SC, Ralph SA, Tilley L. K13, the cytostome, and artemisinin resistance. Trends Parasitol. 2020 Jun;36(6):533–44.

83. Meshnick SR. Artemisinin: mechanisms of action, resistance and toxicity. Int J Parasitol. 2002 Dec 4;32(13):1655–60.

84. Tilley L, Straimer J, Gnädig NF, Ralph SA, Fidock DA. Artemisinin Action and Resistance in Plasmodium falciparum. Trends Parasitol. 2016 Sep;32(9):682–96.

85. Schmidt S, Wichers-Misterek JS, Behrens HM, Birnbaum J, Henshall IG, Dröge J, et al. The Kelch13 compartment contains highly divergent vesicle trafficking proteins in malaria parasites. PLoS Pathog. 2023 Dec;19(12):e1011814.

86. Maier AG, Rug M, O’Neill MT, Brown M, Chakravorty S, Szestak T, et al. Exported proteins required for virulence and rigidity of Plasmodium falciparum-infected human erythrocytes. Cell. 2008 Jul 11;134(1):48–61.

