## Supplementary material for "Single-cell transcriptional changes of artemisinin-sensitive K13^C580^ and artemisinin-resistant K13^580Y^ *Plasmodium falciparum* upon dihydroartemisinin exposure": Supplmental Material

Supplemental Methods…… 2
Supplemental References…… 4
Supplemental Figure S1…… 6
Supplemental Figure S2…… 7
Supplemental Figure S3…… 8
Supplemental Figure S4…… 9
Supplemental Figure S5…… 10
Supplemental Figure S6…… 11
Supplemental Figure S7…… 12
Supplemental Figure S8…… 13
Supplemental Figure S9…… 14
Supplemental Figure S10…… 15
Supplemental Figure S11…… 16
Supplemental Figure S12…… 17
Supplemental Figure S13…… 18
Supplemental Figure S14…… 19
Supplemental Figure S15…… 20
Supplemental Figure S16…… 21
Supplemental Figure S17…… 22
Supplemental Figure S18…… 23
Supplemental Figure S19…… 24
Supplemental Figure S20…… 25
Supplemental Figure S21…… 26
Supplemental Figure S22…… 27
Supplemental Figure S23…… 28
Supplemental Figure S24…… 29
Supplemental Figure S25…… 30
Supplemental Table 1…… 31
Supplemental Table 2…… 32
Supplemental Table 3…… 33
Supplemental Table 4…… 34

**Supplemental Methods**

**SeqWell Library Preparation and Sequencing.** Briefly, 10-15,000 *Pf* infected RBCs from each time point and treatment condition were loaded onto a functionalized polydimethylsiloxane (PDMS) array preloaded with uniquely barcoded mRNA capture beads (Chemgenes; Macosko-2011-10(V+), Wilmington, MA). After cells had settled into wells, the arrays were then sealed with a hydroxylated polycarbonate membrane with a pore size of 10 nm, facilitating buffer exchange while confining biological molecules (such as DNA and RNA) within each well. Following membrane-sealing, subsequent buffer exchange permitted cell lysis, mRNA transcript hybridization to beads, and bead removal before proceeding with reverse transcription. The obtained bead-bound cDNA product then underwent Exonuclease I treatment to remove excess primer before proceeding with second-strand synthesis, then PCR amplification of the cDNA product. The cDNA products were purified using SPRIselect beads (Beckman Coulter, Brea, CA), and were run on a Fragment Analyzer (Agilent Technologies, Santa Clara, CA) to verify the cDNA fragment sizes to be between 0.7-2kbps. The cDNA libraries were then fragmented and amplified for sequencing using the Illumina Nextera XT DNA Library Preparation kit (Illumina, FC-131-1096, San Diego, CA) using custom primers that enabled the specific amplification of only the 3' ends. The libraries were purified, quantified, and then sequenced on a NextSeq550 system using high output 75 cycle kits (Illumina) at an average of 20,000 reads per cell.

**MIP capture, sequencing, and variant calling.** DNA isolates for the isogenic cell lines were captured and sequenced using a drug-resistance MIP panel, which is designed to target mutations and genes associated with antimalarial resistance (1,2). MIP capture and library preparation were performed as previously described (3). Sequencing was conducted using an Illumina NextSeq 550 instrument (150 bp paired-end reads) at Brown University's genomics core (Rhode Island, USA). The MIPtools (v.0.19.12.13; https://github.com/bailey-lab/MIPTools) bioinformatic pipeline was used for processing of sequencing data and variant calling as previously described(1,3,4). The drug-resistance panel included known SNPs in *pfcrt, pfdhfr, pfdhps, pfmdr1, K13,* and other putative drug-resistance genes and has been described elsewhere (1). MIP genotyping results are contained in **Supplemental Table 4**.

**Cell Annotation Pipeline.** Each dataset was processed as follows: the *cell x gene* expression matrix was normalized with SCTransform (5,6) before dimensionality reduction through uniform manifold approximation (UMAP) (7) and clustering through the Leiden algorithm (8). After unsupervised clustering, a first pass was run on our stage annotation pipeline. We calculated pseudobulk gene expression of each cluster, visualizing both literature guided stage markers (9–11) and the top 5 differentially expressed genes per cluster. We correlated our pseudobulk clusters to two bulk RNA sequencing datasets, one a time course experiment measuring RNA transcriptional abundance over the asexual lifecycle (12) and the other focusing on merozoite stages of *P. falciparum (9)*. Next, we mapped our scRNAseq cells to the most recent malaria cell atlas of *P. falciparum (10)* and to a dataset of SmartSeq2 plate based *P. berghei* scRNAseq subsetted to one-to-one orthologs (11). Doublets detected by scrublet and doubletFinder were filtered. Any cells correlated to merozoites were filtered. The remaining cells of all datasets were all integrated with Harmony (13). The stage annotation pipeline was run again, where any cells that were not within the asexual or sexual developmental trajectories were filtered out as doublets (**Supplementary Figure S8** and **S9**). After those doublets were filtered, the stage annotation pipeline was run for a final time to result in the final stage annotations (**Supplemental Figure S10**-**S12**). The proportions of each stage are shown in **Supplemental Figure S13**. Plots generated after each step in this pipeline are presented in an interactive HTML at https://github.com/sconnelly007/K13_mt_v_wt_sc/clustering_summary.html.

**Fine Grained Differential Expression with MELD**. The normalized cell x gene expression matrix for each strain was exported from Seurat and imported as an annotated data object (14) in Python (v3.9.15). Only the cells from the DMSO or DHA treatment groups were used for this analysis. Each strain was analyzed separately, and PCA was performed on each gene expression matrix. The top 100 PCs from the PCA embedding were used to create a Potential of Heat diffusion for Affinity-based Transition Embedding (PHATE) embedding, with default settings (15). MELD was fit to the normalized cell x gene expression matrix over a range of k-nearest neighbors (KNNs) and beta, a smoothing parameter, values. During fitting, a random probability density function (PDF) is generated over the data, and cells are randomly assigned to DHA or DMSO treatment conditions. MELD was then fit to the normalized cell x gene matrix with the given parameters, and the Mean Square Error (MSE) of the DHA likelihoods between the DHA condition and the random PDF was calculated (see **Supplemental Figure S15**). This process was repeated 25 times for each KNN, over the range of beta values. The average MSE was calculated, and the parameters with the lowest mean MSE (shown as a red dot in **Supplemental Figure S16**) were selected. The optimal parameters for each strain were then used downstream for running MELD.

For each strain, MELD was run per time point, with the optimal parameters per strain, the timepoint's PCA embedding, and treatment condition (DMSO or DHA) labels as input, similar to previous work (16). The output was a continuous score of experimental perturbation, referred to as the DHA likelihood. For each malaria cell cycle stage, the PCA embedding, treatment labels, and DHA likelihoods were used as input for vertex frequency clustering (VFC). This PCA embedding was then used by PHATE to visualize the VFC results. VFC results are in **Supplemental Figures S18-S22**.

Differential expression between the VFC clusters with the highest DHA likelihood per timepoint was performed with the MAST (17) package. All differential expression results are present in **Supplemental Table S1**. For visualization in **Figures 1-3**, these genes were pruned to the top 5 differentially expressed hypervariable genes by absolute log2FC. Then, annotated genes were selected and ranked by log2FC to find the top 50 genes by absolute log2FC. The vectors of the top 50 (**Figure 1**) or top 30 (**Figures 2** and **3**) genes were concatenated and filtered to unique genes.

**Supplemental References**


1. Aydemir O, Janko M, Hathaway NJ, Verity R, Mwandagalirwa MK, Tshefu AK, et al. Drug-resistance and population structure of Plasmodium falciparum across the Democratic Republic of Congo using high-throughput molecular inversion probes. J Infect Dis. 2018 Aug 14;218(6):946–55.

2. Verity R, Aydemir O, Brazeau NF, Watson OJ, Hathaway NJ, Mwandagalirwa MK, et al. The impact of antimalarial resistance on the genetic structure of Plasmodium falciparum in the DRC. Nat Commun. 2020 Apr 30;11(1):2107.

3. Moser KA, Madebe RA, Aydemir O, Chiduo MG, Mandara CI, Rumisha SF, et al. Describing the current status of Plasmodium falciparum population structure and drug resistance within mainland Tanzania using molecular inversion probes. Mol Ecol. 2021 Jan;30(1):100–13.

4. Fola AA, Feleke SM, Mohammed H, Brhane BG, Hennelly CM, Assefa A, et al. Plasmodium falciparum resistant to artemisinin and diagnostics have emerged in Ethiopia. Nat Microbiol. 2023 Oct;8(10):1911–9.

5. Hafemeister C, Satija R. Normalization and variance stabilization of single-cell RNA-seq data using regularized negative binomial regression. Genome Biol. 2019 Dec 23;20(1):296.

6. Choudhary S, Satija R. Comparison and evaluation of statistical error models for scRNA-seq. Genome Biol. 2022 Jan 18;23(1):27.

7. Becht E, McInnes L, Healy J, Dutertre CA, Kwok IWH, Ng LG, et al. Dimensionality reduction for visualizing single-cell data using UMAP. Nat Biotechnol [Internet]. 2018 Dec 3; Available from: http://dx.doi.org/10.1038/nbt.4314

8. Traag VA, Waltman L, van Eck NJ. From Louvain to Leiden: guaranteeing well-connected communities. Sci Rep. 2019 Mar 26;9(1):5233.

9. Reers AB, Bautista R, McLellan J, Morales B, Garza R, Bol S, et al. Histone modification analysis reveals common regulators of gene expression in liver and blood stage merozoites of Plasmodium parasites. Epigenetics Chromatin. 2023 Jun 15;16(1):25.

10. Dogga SK, Rop JC, Cudini J, Farr E, Dara A, Ouologuem D, et al. A single cell atlas of sexual development in Plasmodium falciparum. Science. 2024 May 3;384(6695):eadj4088.

11. Howick VM, Russell AJC, Andrews T, Heaton H, Reid AJ, Natarajan K, et al. The Malaria Cell Atlas: Single parasite transcriptomes across the complete Plasmodium life cycle. Science [Internet]. 2019 Aug 23;365(6455). Available from: http://dx.doi.org/10.1126/science.aaw2619

12. Painter HJ, Chung NC, Sebastian A, Albert I, Storey JD, Llinás M. Genome-wide real-time in vivo transcriptional dynamics during Plasmodium falciparum blood-stage development. Nat Commun [Internet]. 2018 Jul 9;9(1). Available from: http://dx.doi.org/10.1038/s41467-018-04966-3

13. Korsunsky I, Millard N, Fan J, Slowikowski K, Zhang F, Wei K, et al. Fast, sensitive and accurate integration of single-cell data with Harmony. Nat Methods. 2019 Dec;16(12):1289–96.

14. Wolf FA, Angerer P, Theis FJ. SCANPY: large-scale single-cell gene expression data analysis. Genome Biol. 2018 Feb 6;19(1):15.

15. Moon KR, van Dijk D, Wang Z, Gigante S, Burkhardt DB, Chen WS, et al. Visualizing structure and transitions in high-dimensional biological data. Nat Biotechnol. 2019 Dec;37(12):1482–92.

16. Burkhardt DB, Stanley JS 3rd, Tong A, Perdigoto AL, Gigante SA, Herold KC, et al. Quantifying the effect of experimental perturbations at single-cell resolution. Nat Biotechnol. 2021 May;39(5):619–29.

17. Finak G, McDavid A, Yajima M, Deng J, Gersuk V, Shalek AK, et al. MAST: a flexible statistical framework for assessing transcriptional changes and characterizing heterogeneity in single-cell RNA sequencing data. Genome Biol. 2015 Dec 10;16:278.

18. Neavin D, Senabouth A, Arora H, Lee JTH, Ripoll-Cladellas A, sc-eQTLGen Consortium, et al. Demuxafy: improvement in droplet assignment by integrating multiple single-cell demultiplexing and doublet detection methods. Genome Biol. 2024 Apr 15;25(1):94.

19. Street K, Risso D, Fletcher RB, Das D, Ngai J, Yosef N, et al. Slingshot: cell lineage and pseudotime inference for single-cell transcriptomics. BMC Genomics [Internet]. 2018 Dec;19(1). Available from: http://dx.doi.org/10.1186/s12864-018-4772-0

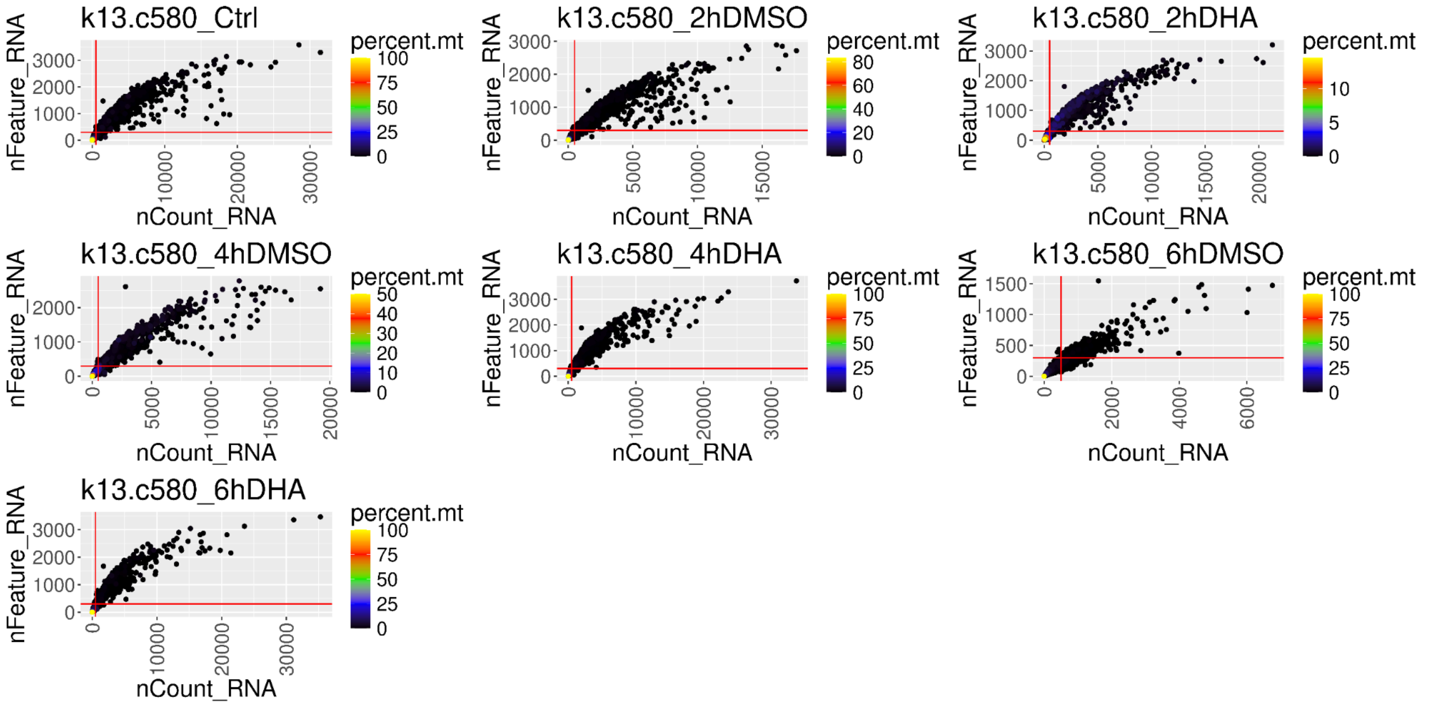


**Supplementary Figure S1 - Quality Control Filtering of K13$^{C580}$ cells.** Cells are plotted by the number of UMI counts (nCount_RNA) vs. the number of genes (nFeature_RNA), while being colored by the percent mitochondrial gene expression (percent.mt). Filtering thresholds are shown in red (greater than 500 UMIs for nCount_RNA and greater than 300 genes for nFeature_RNA). Additionally, cells with less than 20% mitochondrial gene expression were retained.

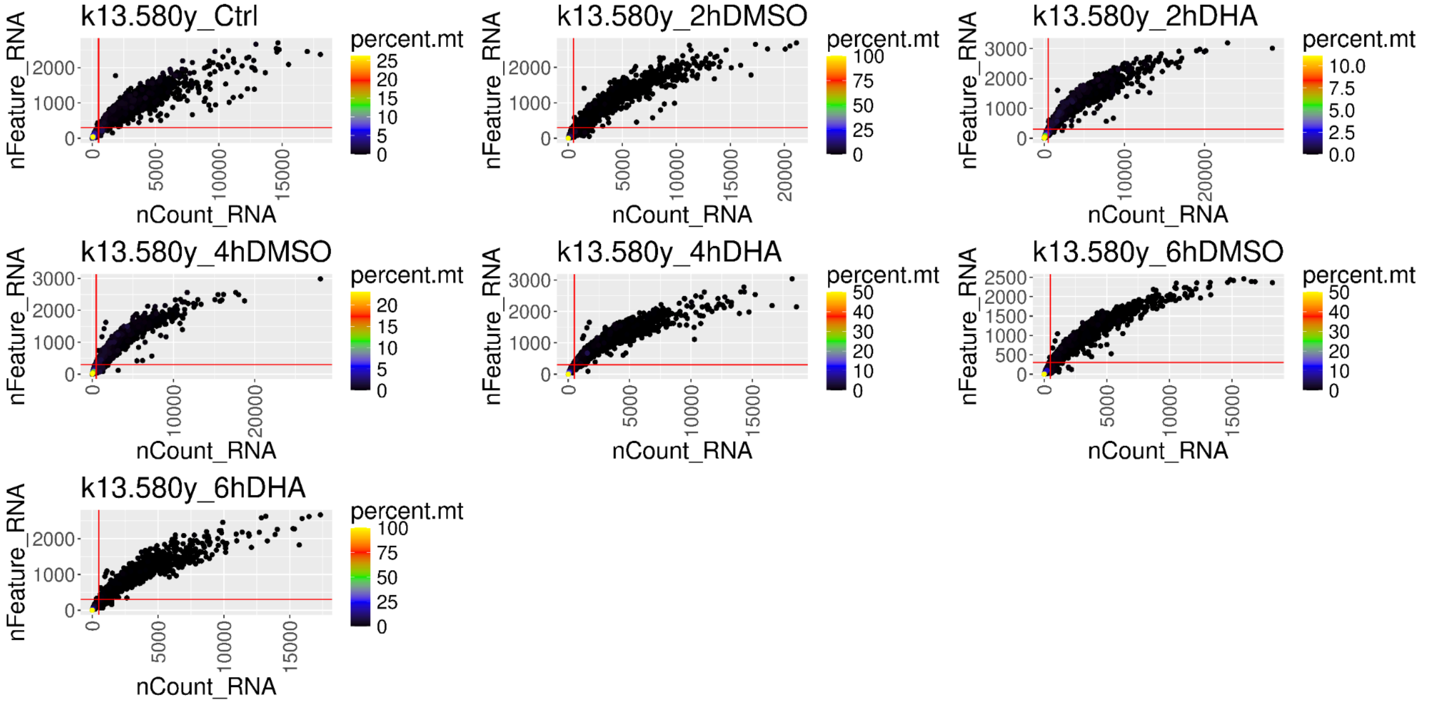


**Supplementary Figure S2 - Quality Control Filtering of K13$^{580Y}$ cells.** Cells are plotted by the number of UMI counts (nCount_RNA) vs. the number of genes (nFeature_RNA), while being colored by the percent mitochondrial gene expression (percent.mt). Filtering thresholds are shown in red (greater than 500 UMIs for nCount_RNA and greater than 300 genes for nFeature_RNA). Additionally, cells with less than 20% mitochondrial gene expression were retained.

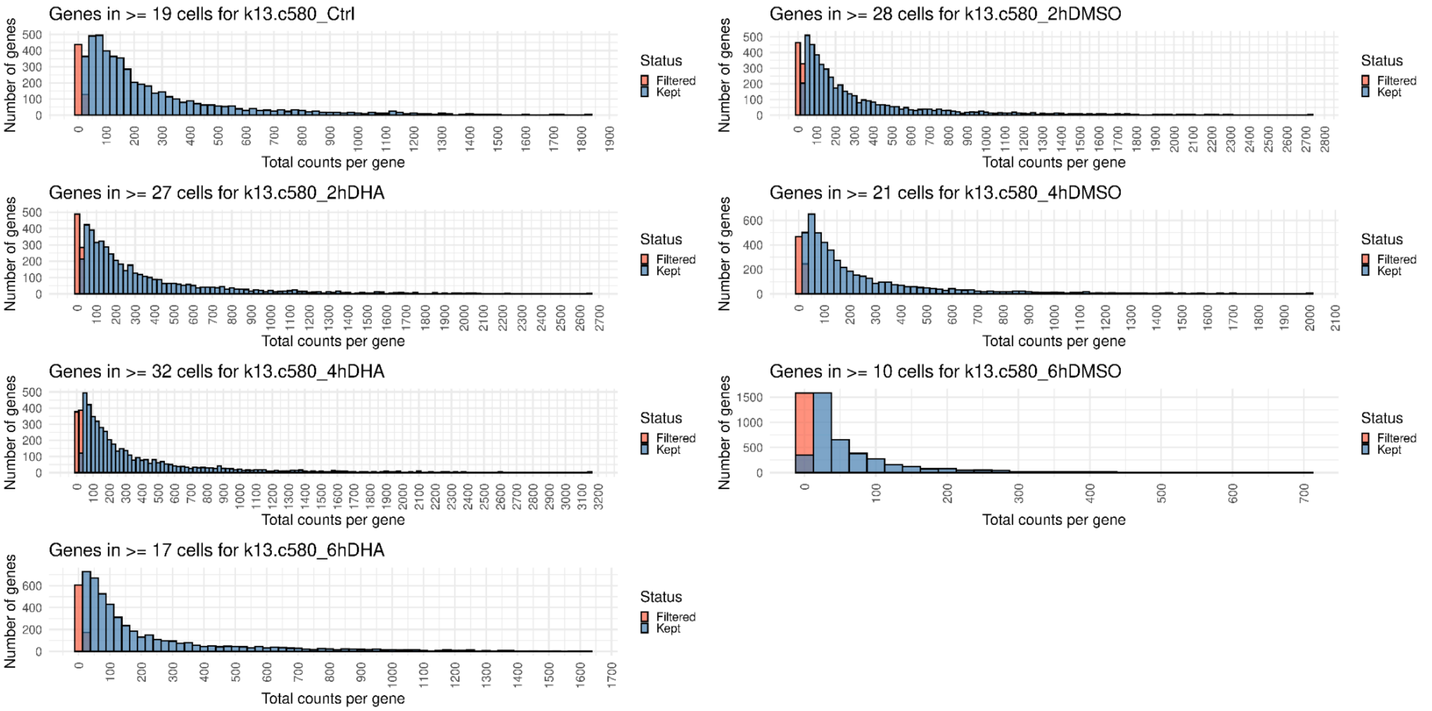


**Supplementary Figure S3 - Quality Control Filtering of Genes in the K13[C580] Dataset.** Each plot shows a histogram of genes binned by the number of UMI counts over all cells. Genes were retained with a threshold determined by the maximum between two options; if they were expressed in at least 10 cells or expressed in at least 1% of all cells.

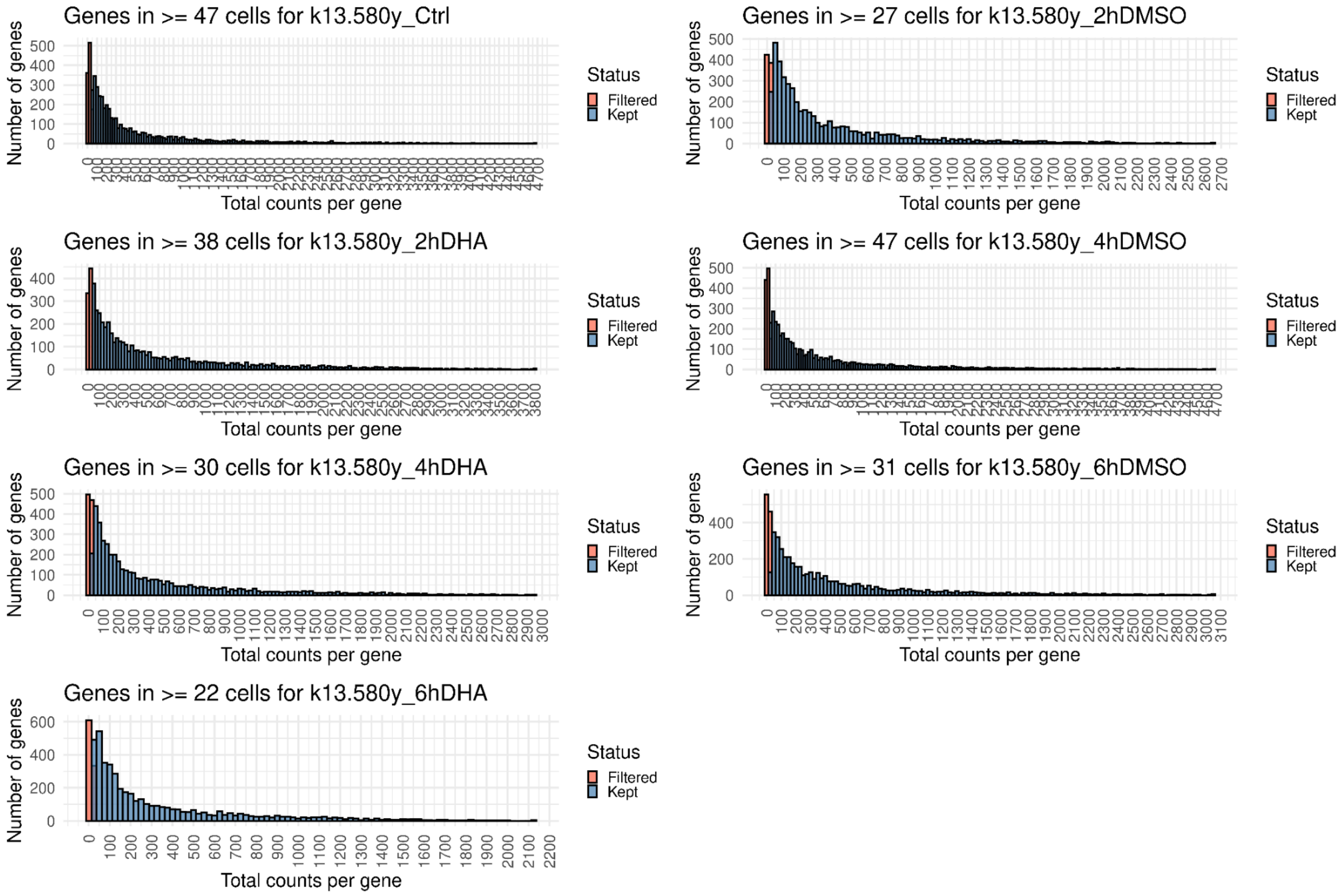


**Supplementary Figure S4 - Quality Control Filtering of Genes in the K13$^{580Y}$ Dataset.** Each plot shows a histogram of genes binned by the number of UMI counts over all cells. Genes were retained with a threshold determined by the maximum between two options; if they were expressed in at least 10 cells or expressed in at least 1% of all cells.

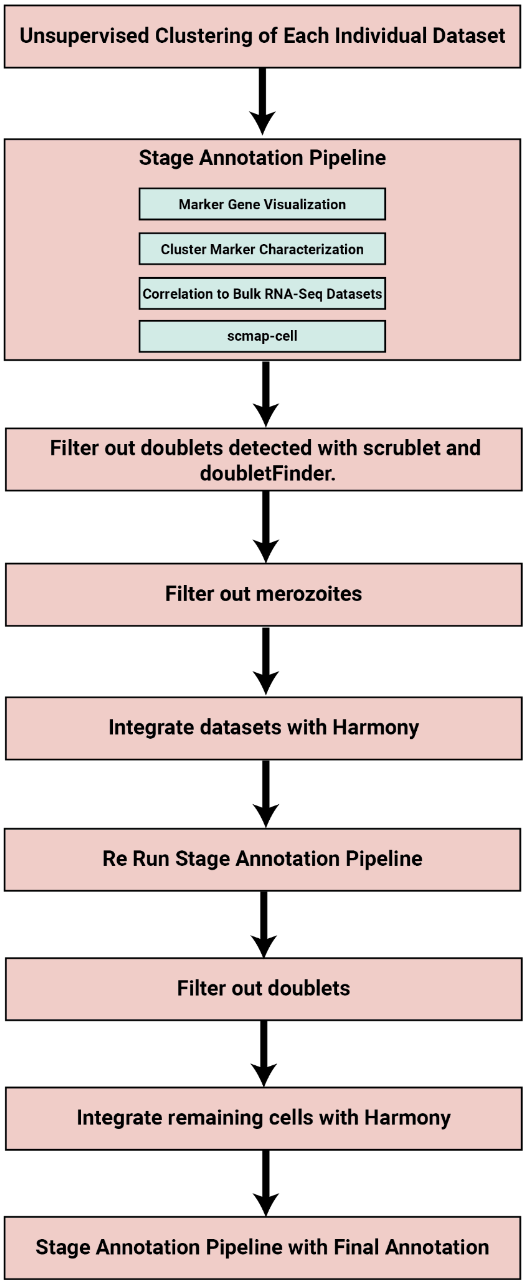


**Supplementary Figure S5 - Cell Annotation Pipeline.** A flowchart of the cell annotation pipeline is detailed alongside our Supplementary Methods section above.

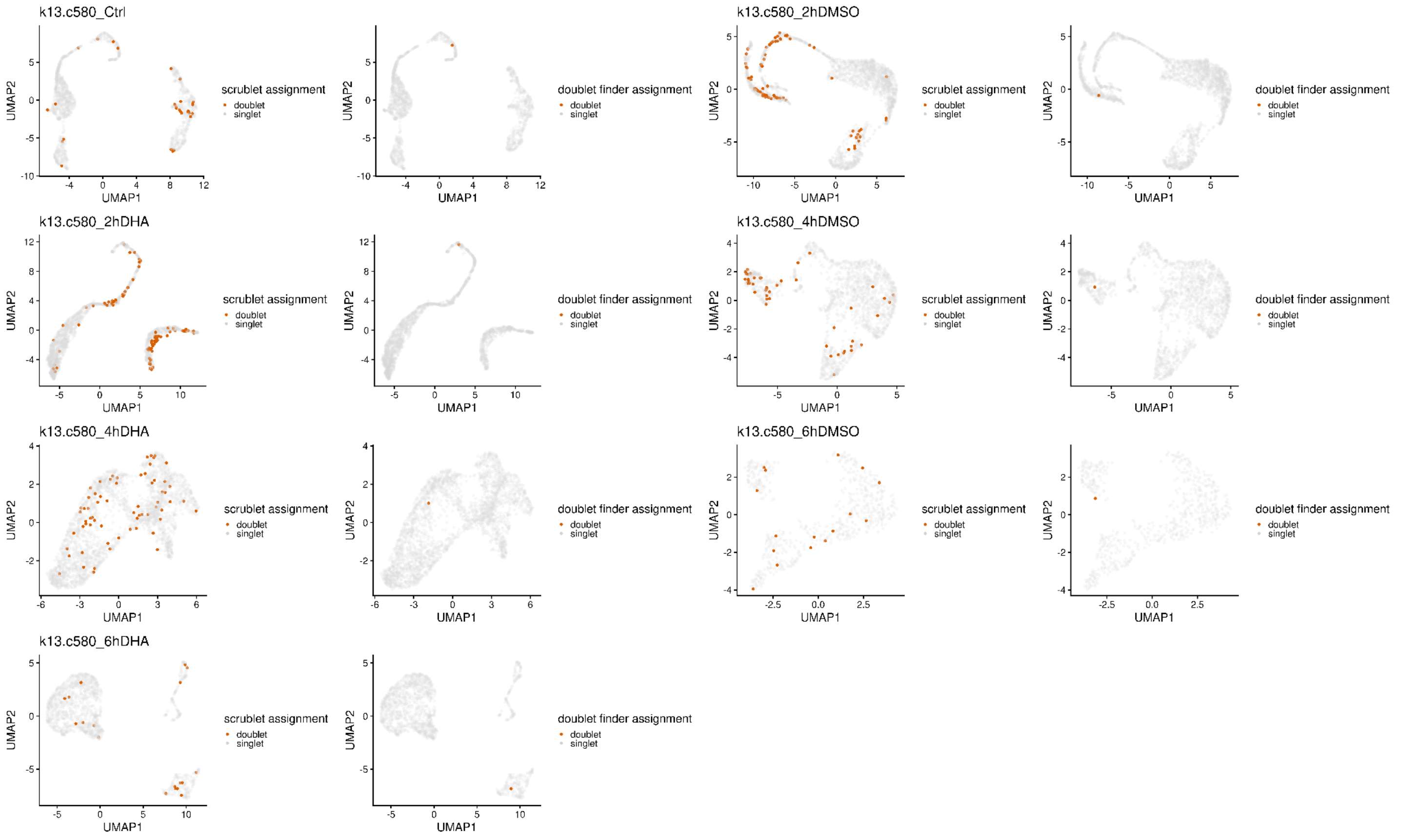


**Supplementary Figure S6 - Doublets Detected in the K13$^{C580}$ Dataset.** The number of doublets was estimated to be 2.37% based on previous SeqWell publications. This value was input into both scrublet and doubletFinder to simulate doublets. We utilized 0.13 as the doublet detection threshold in scrublet and the implementation of doubletFinder in the Demuxafy (18) package to simulate doublets.

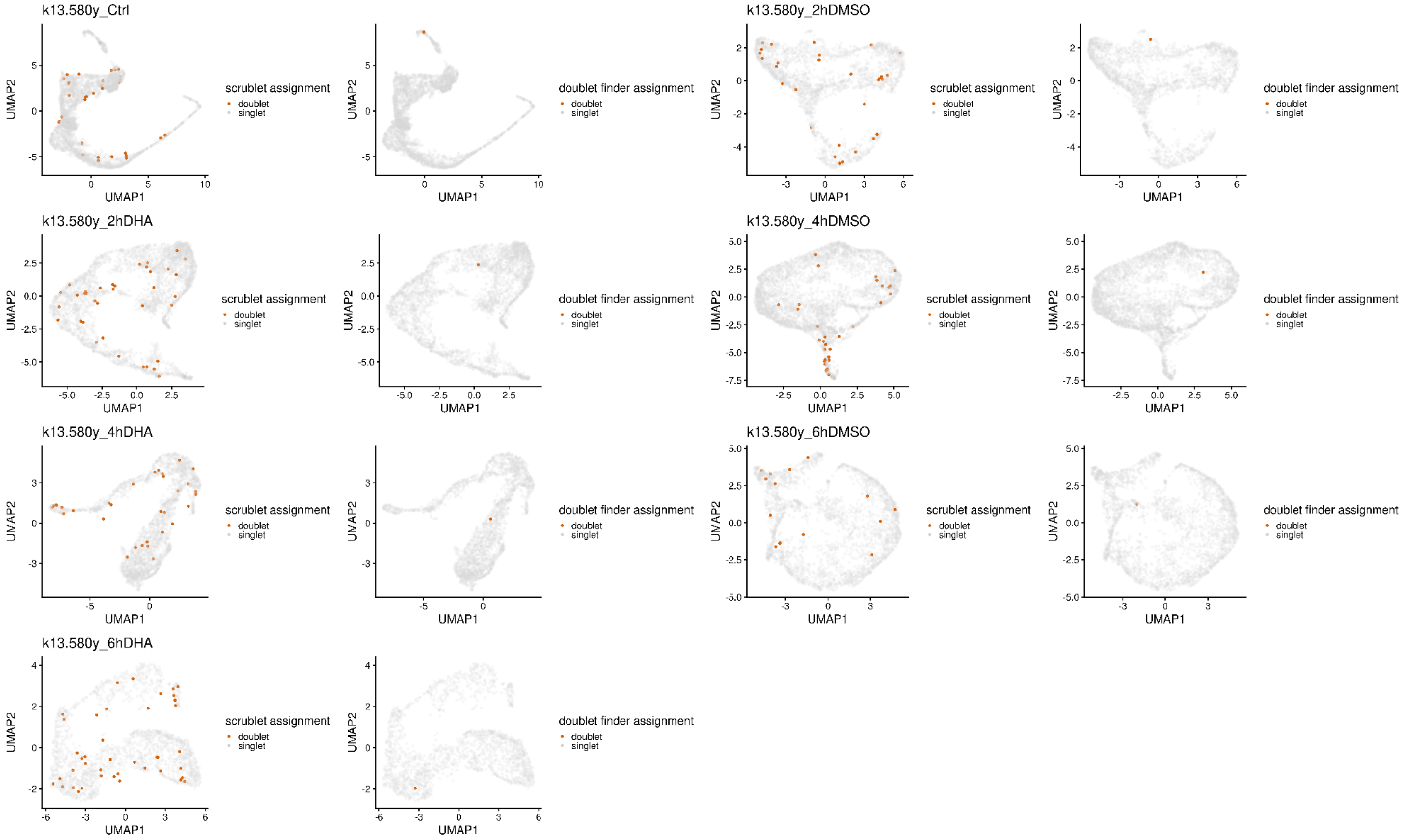


**Supplementary Figure S7 - Doublets Detected in the $K13^{580Y}$ Dataset.** The number of doublets was estimated to be 2.37% based on previous SeqWell publications. This value was input into both scrublet and doubletFinder to simulate doublets. We utilized 0.13 as the doublet detection threshold in scrublet and the implementation of doubletFinder in the Demuxafy (18) package to simulate doublets.

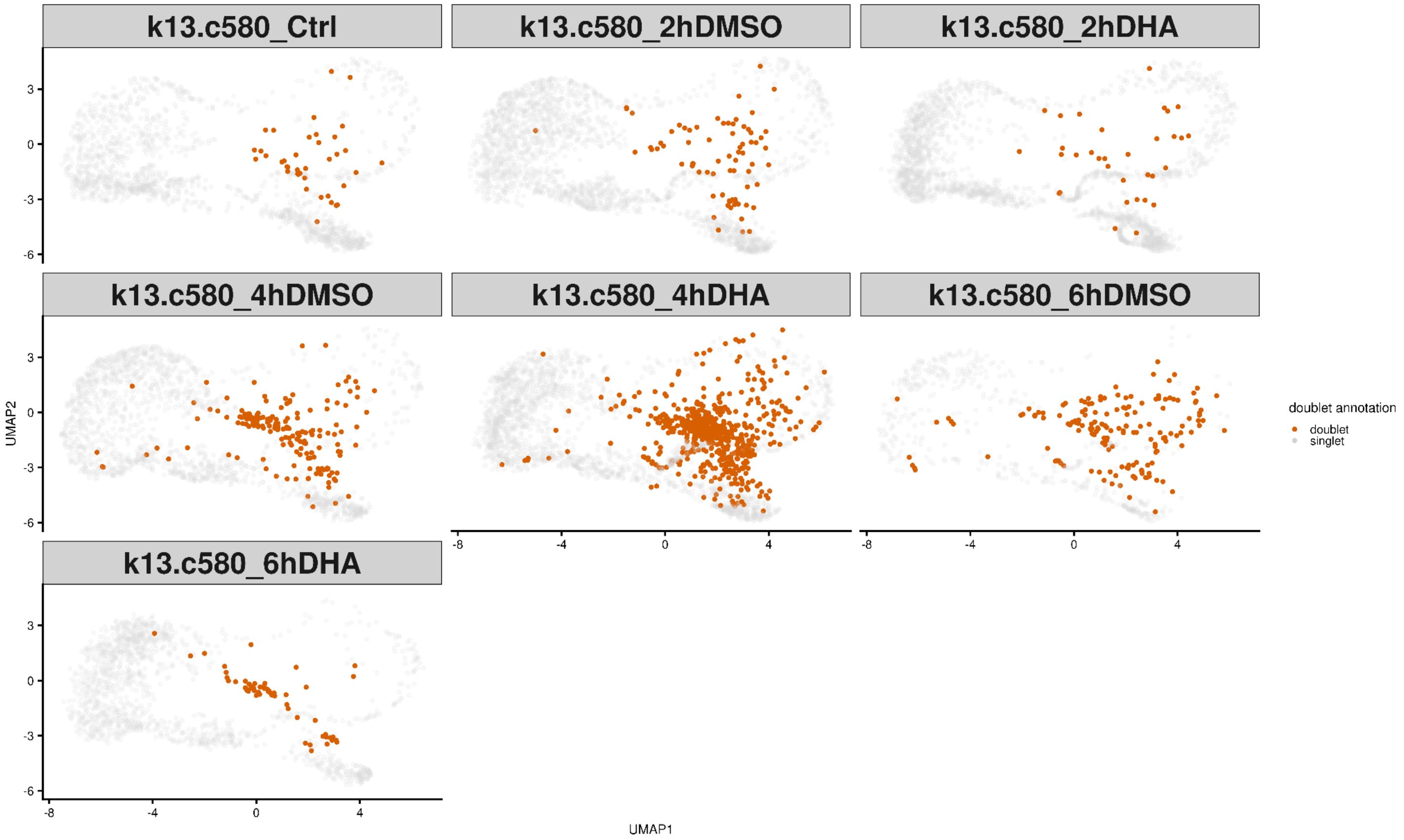


**Supplementary Figure S8 - Doublets Detected after Integrating in K13$^{C580}$.** After integration of the K13$^{C580}$ and K13$^{580Y}$ datasets, doublets were labeled as those cells embedded outside of the asexual and sexual lifecycle trajectories.

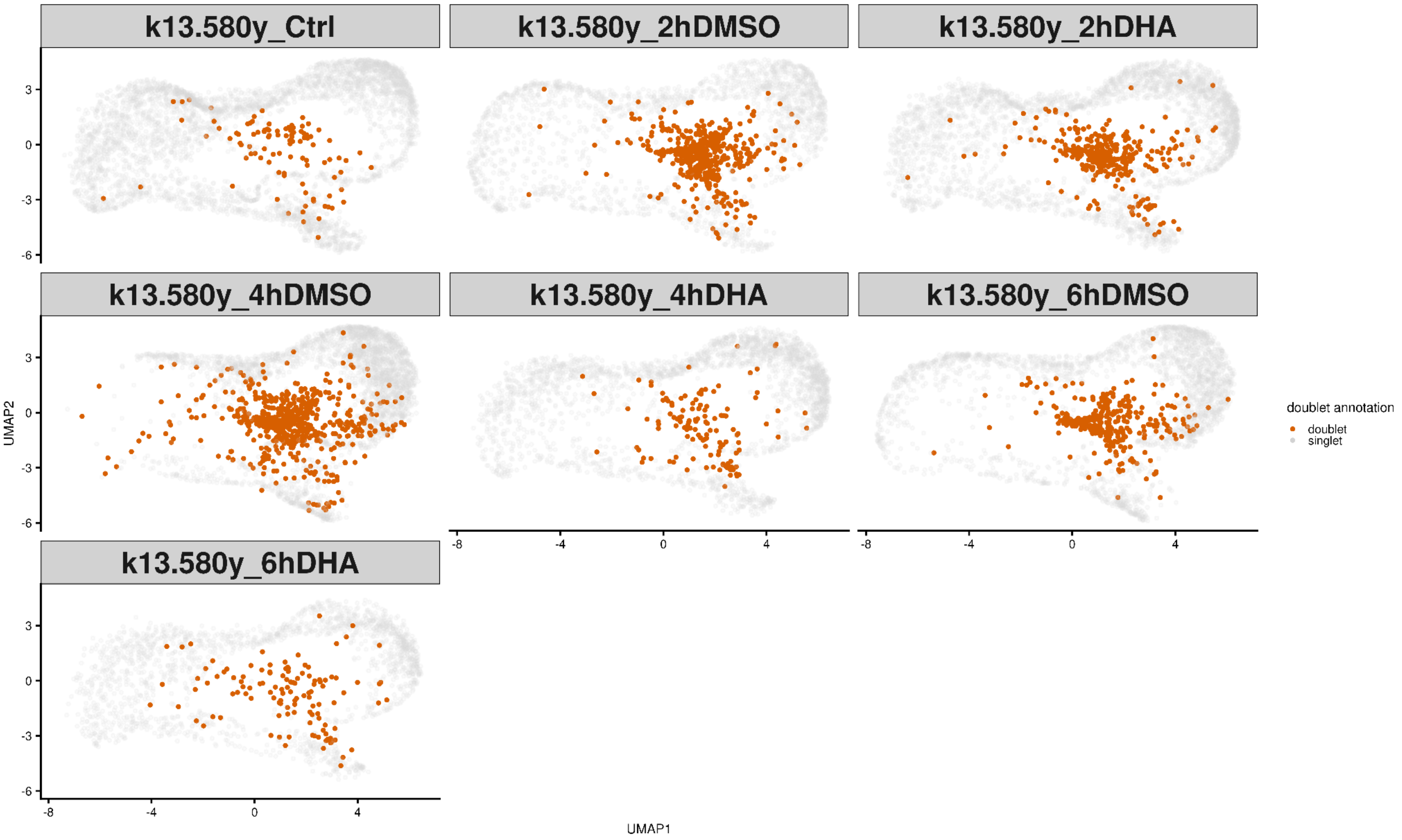


**Supplementary Figure S9 - Doublets Detected after Integrating in K13$^{580Y}$.** After integration of the K13$^{C580}$ and K13$^{580Y}$ datasets, doublets were labeled as those cells embedded outside of the asexual and sexual lifecycle trajectories.

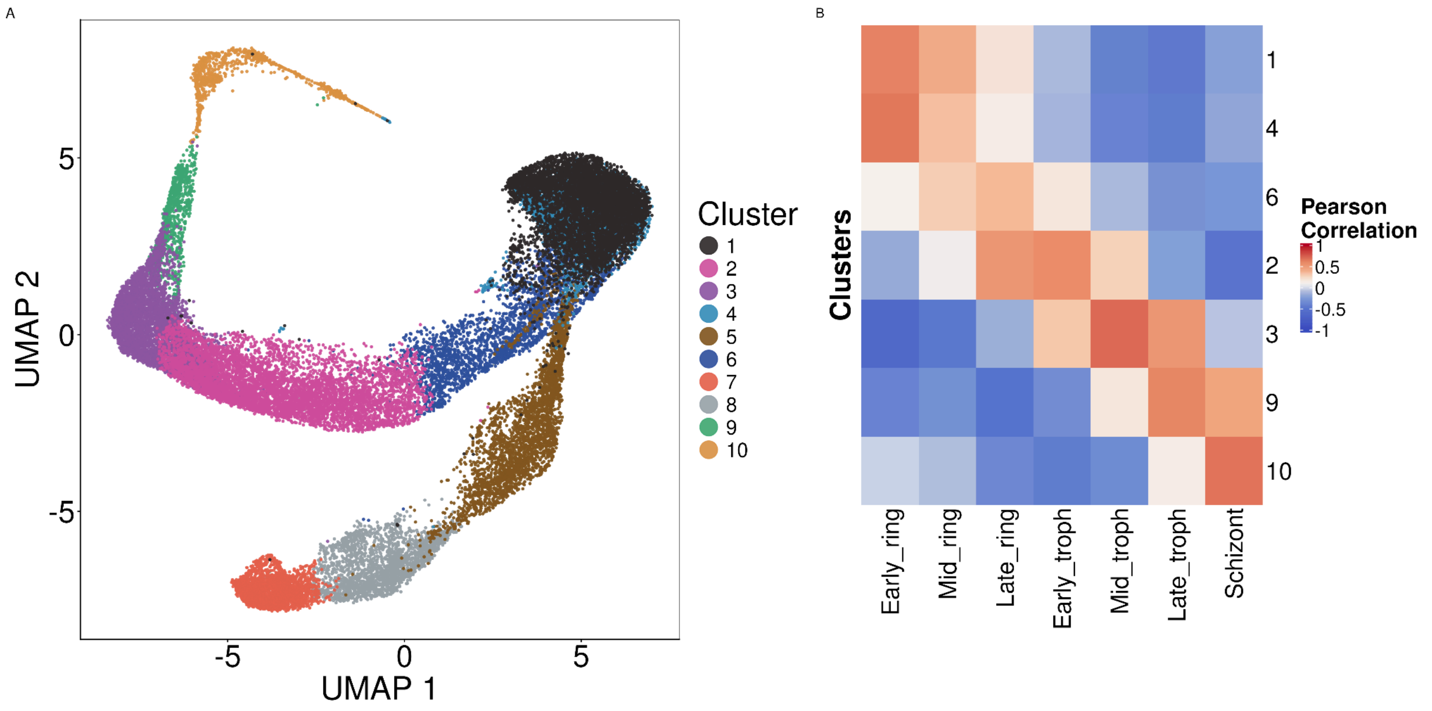


**Supplementary Figure S10 - Unsupervised Clustering and Correlation to Painter *et al*..** A) After filtering out all doublets and merozoites, unsupervised clustering was performed over the entire integrated dataset. B) The correlation results between the clusters annotated as asexual stages and the Painter *et al.* dataset. Other clusters were annotated based on marker genes (cluster 5 as sexually committed rings (Supplementary Figure S11) and clusters 7 and 8 as gametocytes (Supplementary Figure S12).

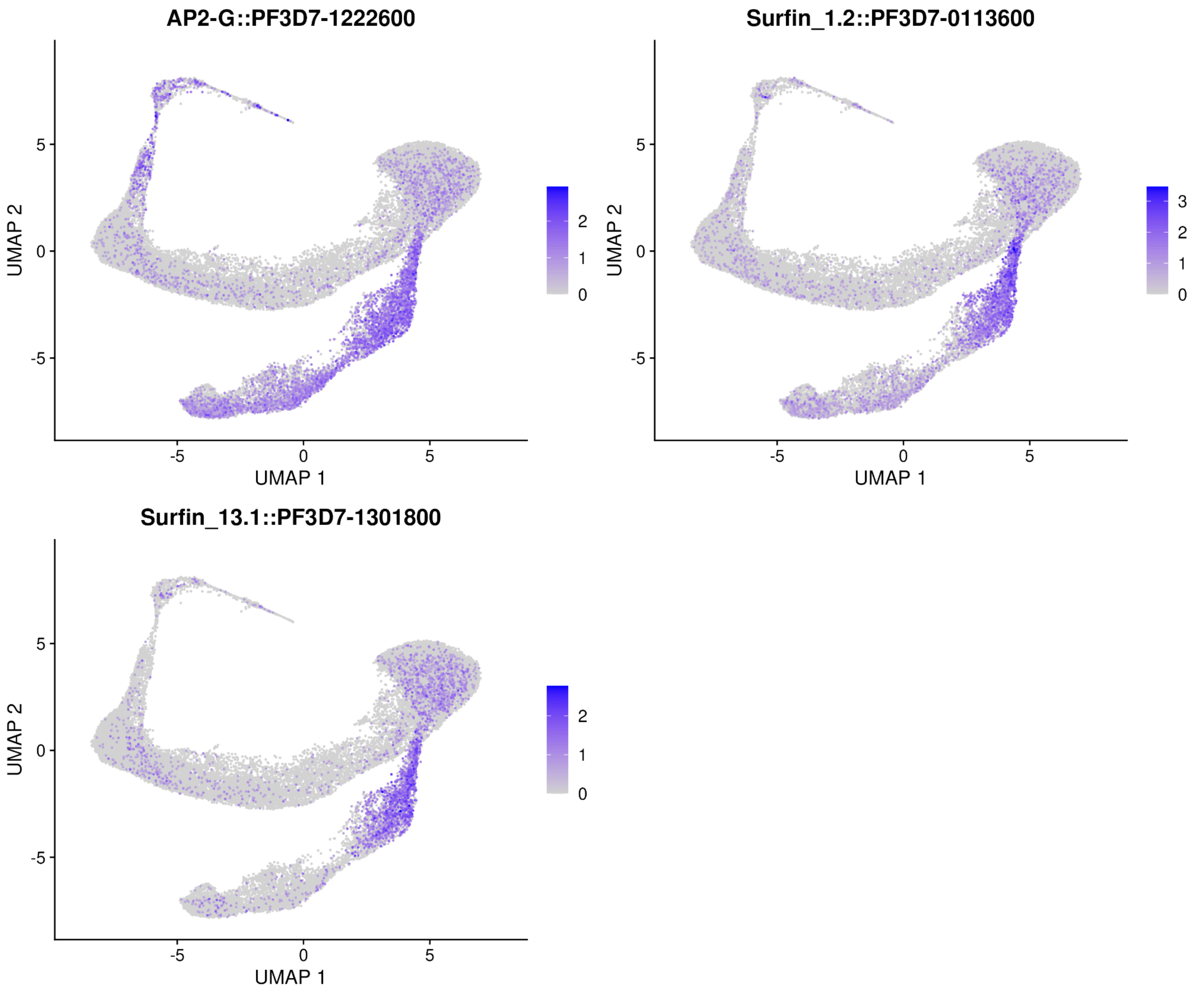


**Supplementary Figure S11 - Expression of Sexually Committed Ring Markers Over Integrated Dataset.** Expression of sexually committed ring markers (*AP2-G*, *Surfin 1.2* and *Surfin 13.1*) from Prajapati *et al.* 2020 are shown over the integrated dataset's UMAP. The expression is concentration in Cluster 5.

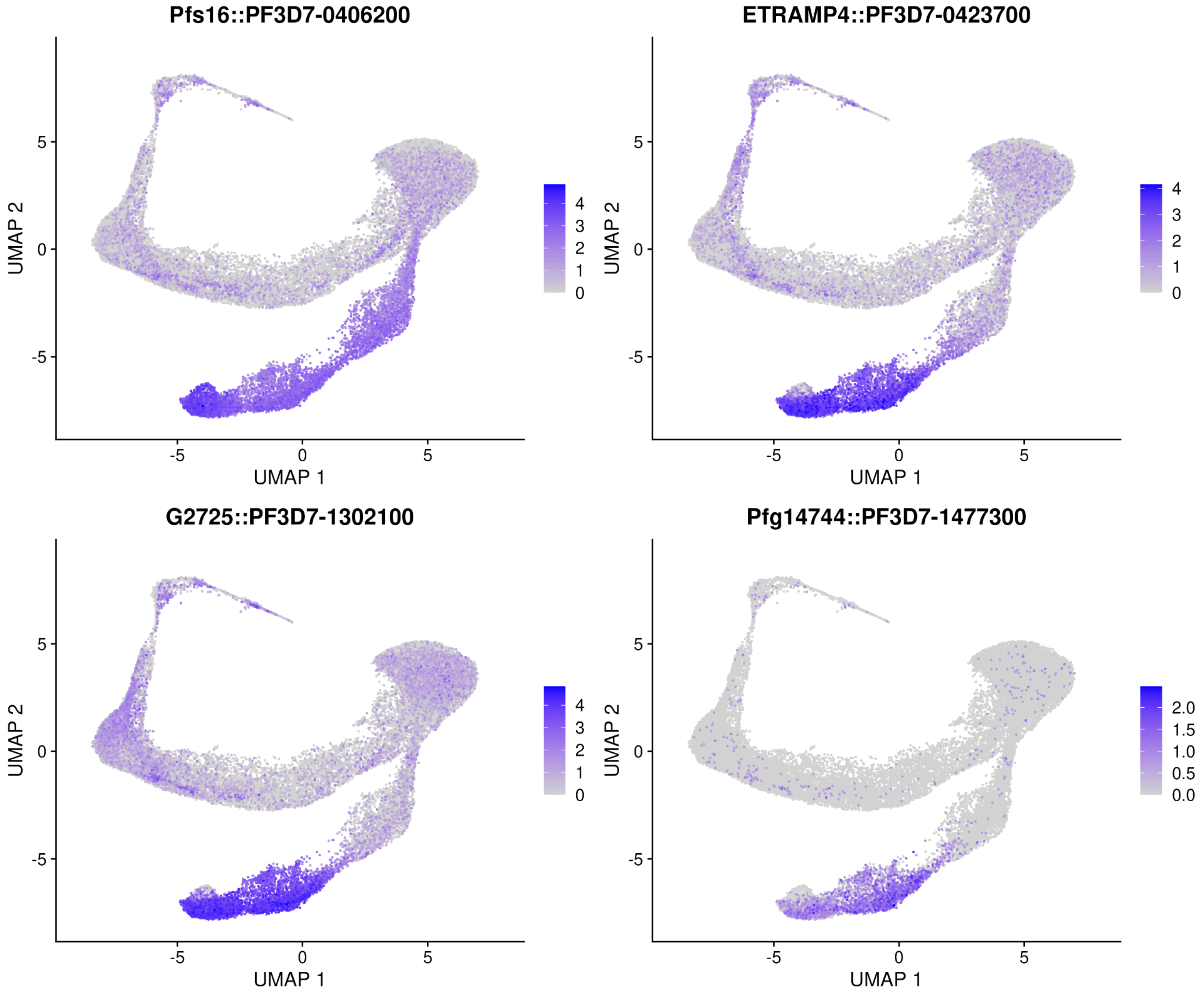


**Supplementary Figure S12 - Expression of Gametocyte Markers Over Integrated Dataset.** Expression of gametocyte markers from Dogga *et al.* are plotted over the entire integrated dataset. Clusters 7 and 8 show the highest expression of these markers.

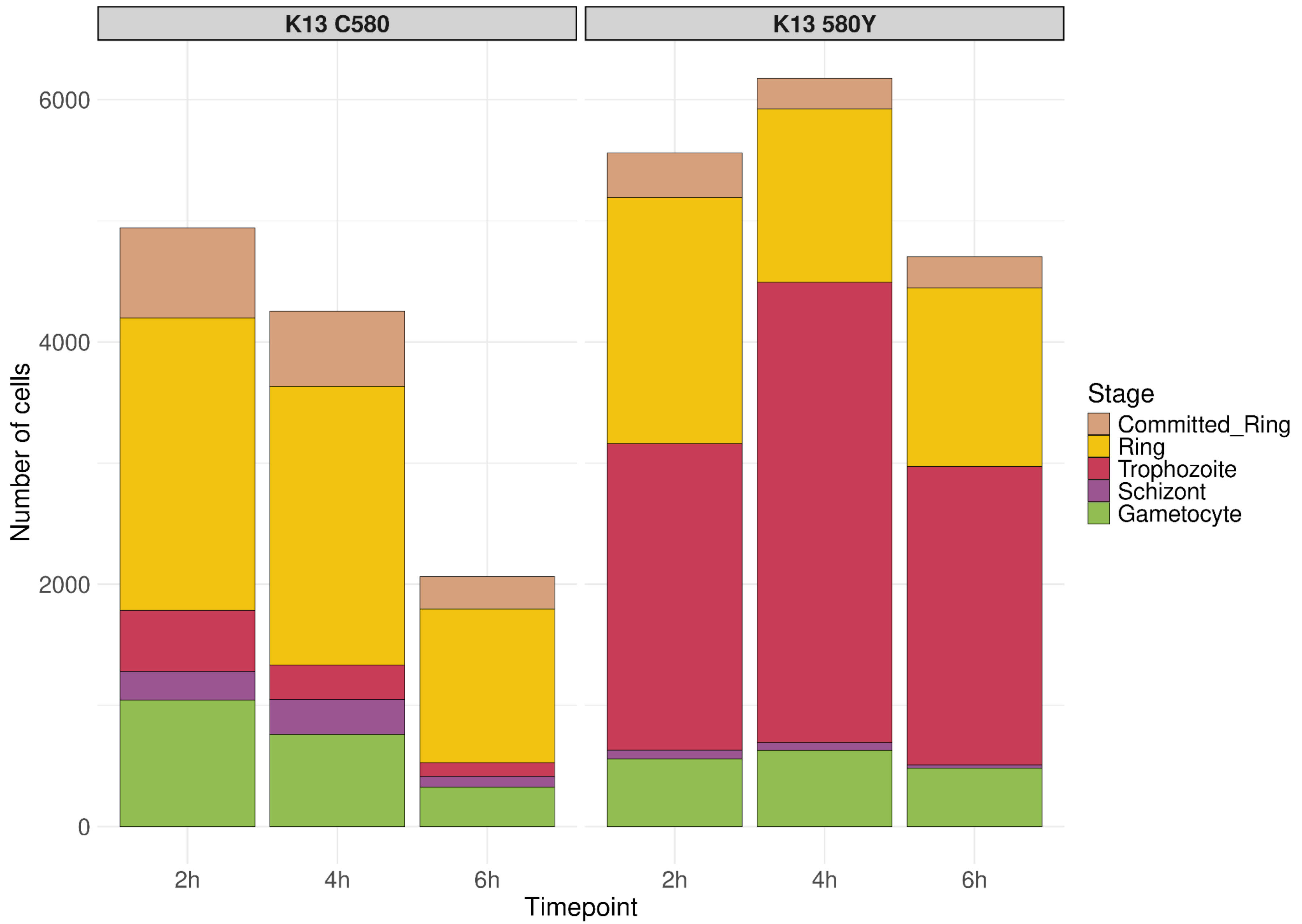


**Supplementary Figure S13 - Stage Distribution Per Strain Over Each Timepoint.** For each strain and timepoint, a stacked bar plot is plotted that shows the number of each parasite stage.

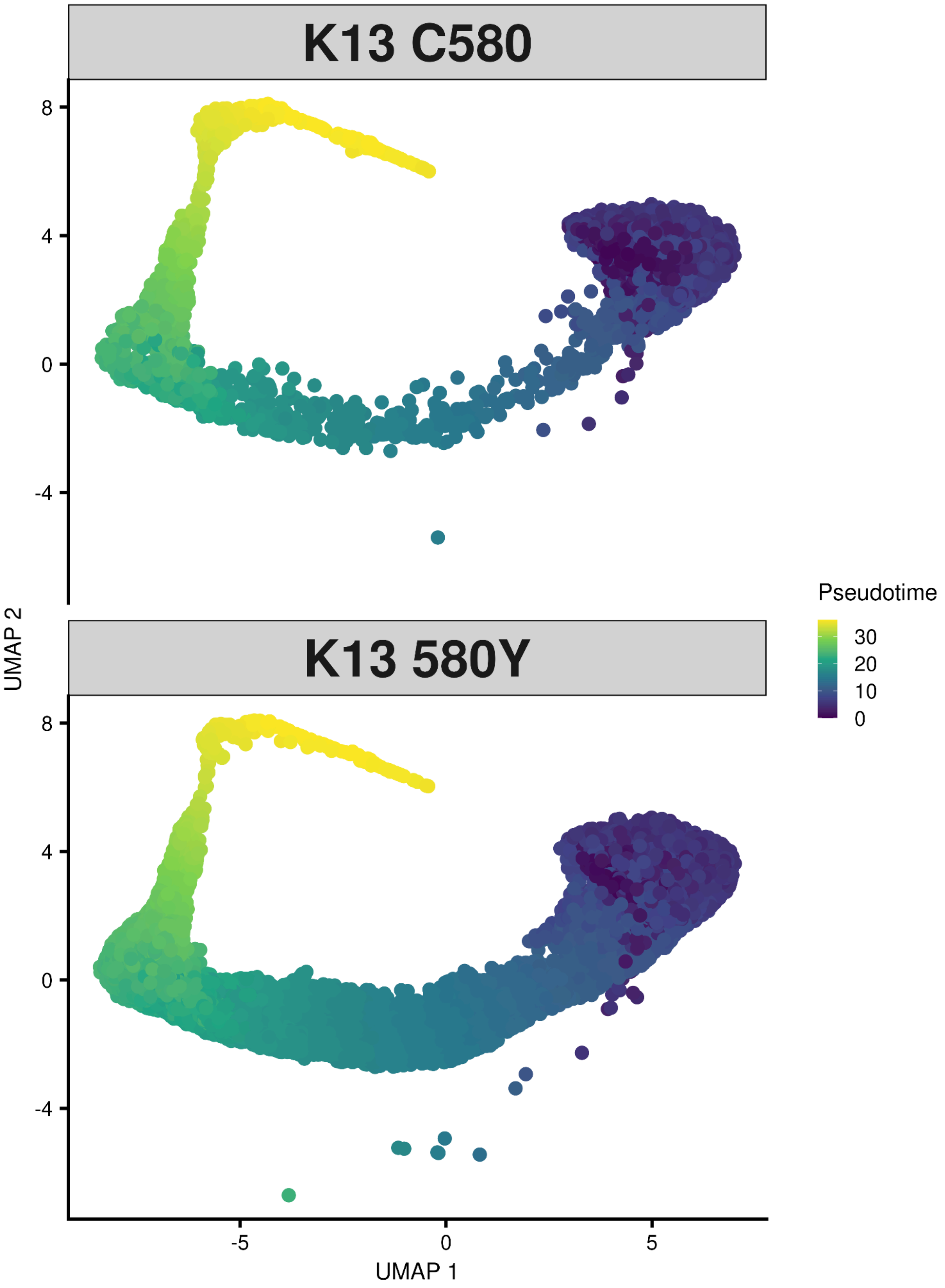


**Supplemental Figure S14. Pseudotime over the asexual cycle, separated by strain.** Pseudotime was calculated using slingshot for each strain (top: $K13^{C580}$, bottom: $K13^{580Y}$) over the ring, trophozoite, and schizont stages. Each cell is assigned a pseudotime value. Lower values correspond to earlier developmental pseudotime (ring stage) and are colored purple. Pseudotime values then increase through the trophozoite stage (green) and the schizont stage (yellow).

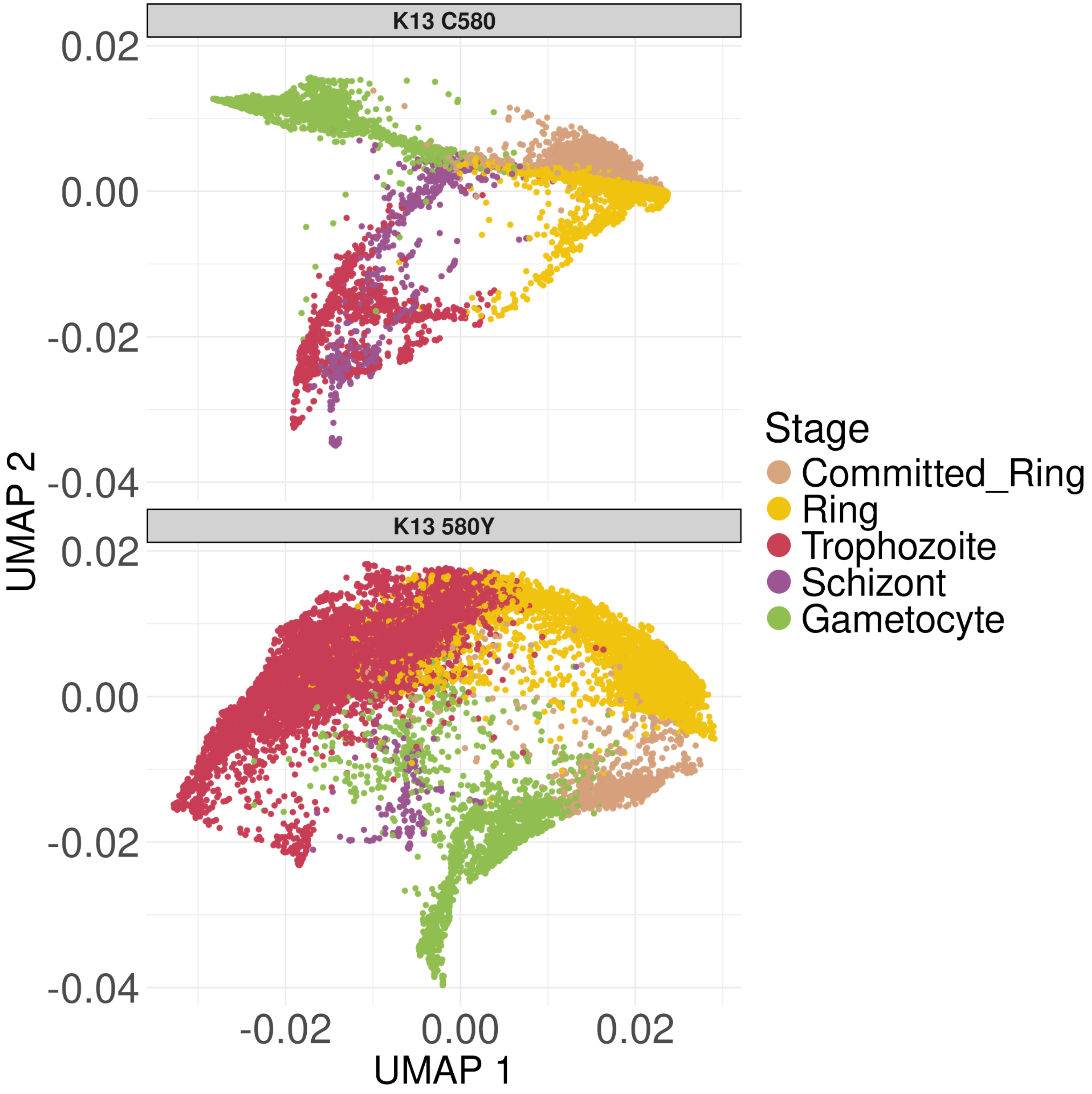


**Supplemental Figure S15. PHATE Embedding.** Normalized counts exported from the integrated Seurat object were exported and used to create a PHATE embedding for each strain (top: K13$^{C580}$, bottom: K13$^{580Y}$). Cells are colored by stage annotation.

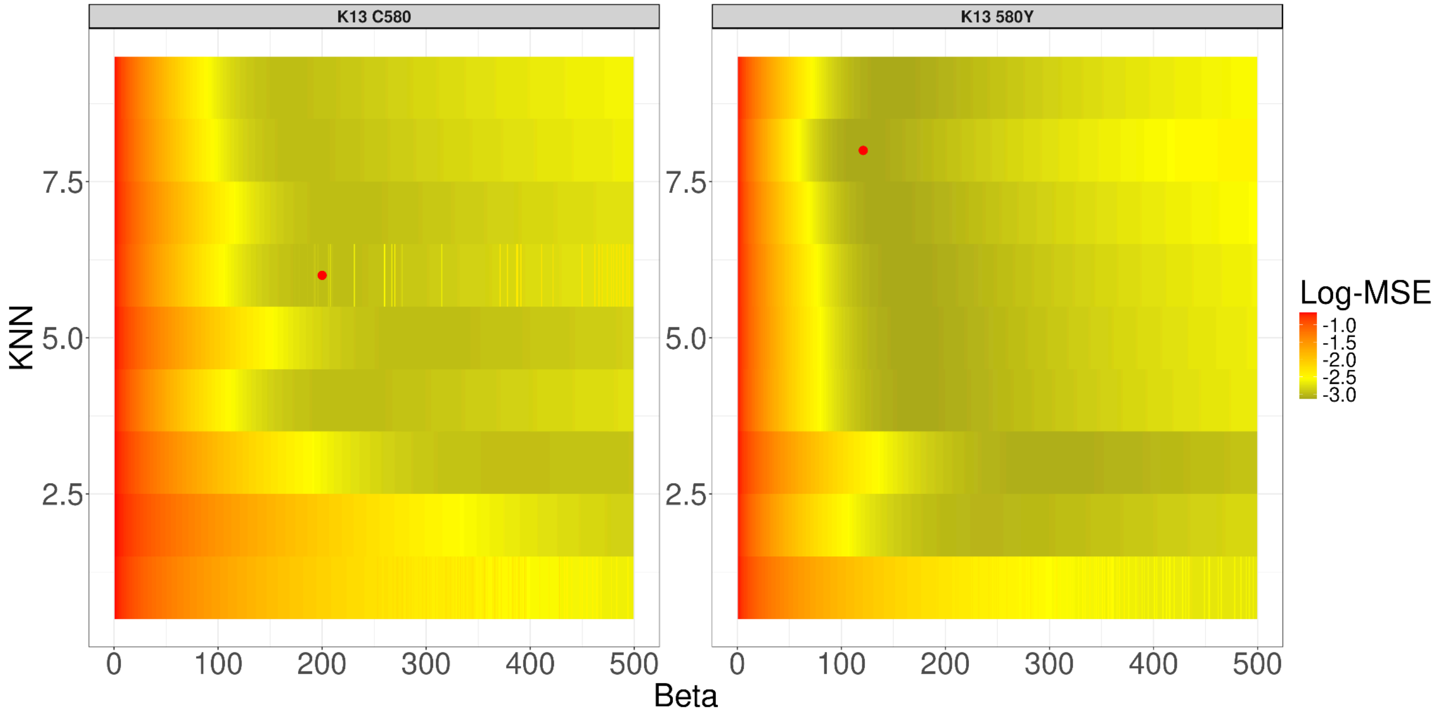


**Supplemental Figure S16. Optimization of MELD algorithm over different KNN (y axis) and beta (x axis) parameters.** As described in the Supplemental Methods, MELD was fit to each strain's normalized *cell x gene* matrix separately and over a range of KNN and beta values. The red dot on each plot is the optimal parameters used to run MELD for each line.

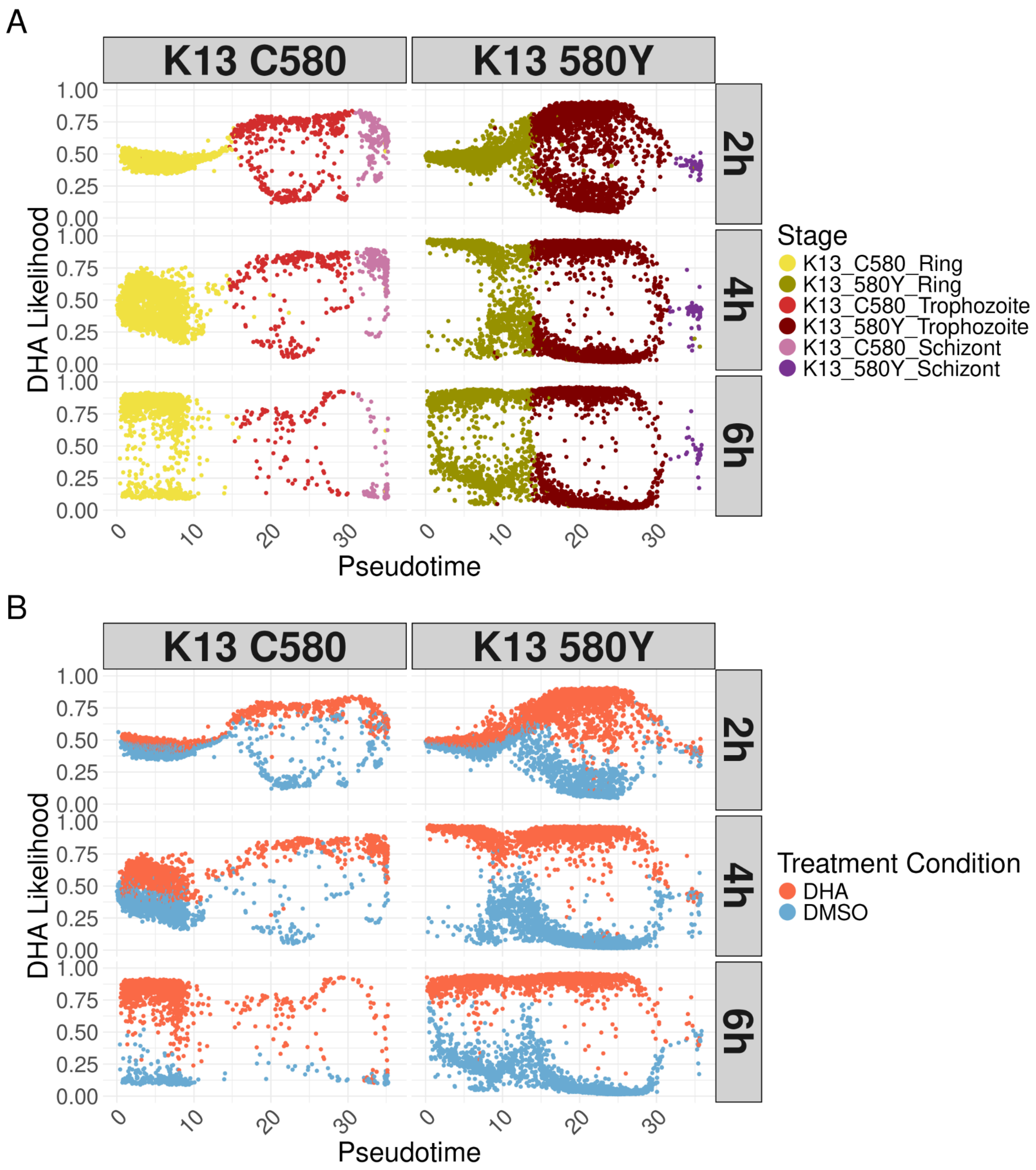


**Supplemental Figure S17. DHA treated parasites have higher DHA likelihoods over the lifecycle than DMSO treated parasites.** Pseudotime, calculated per strain through slingshot (19), is plotted on the x-axis while the DHA likelihood output from MELD is plotted on the y-axis for each facet of the figure. Cells are colored by stage annotation (A) or treatment condition (B). The left column plots K13$^{C580}$ cells while the right column plots K13$^{580Y}$ cells. Each row is a different timepoint, with the first row being the 2 hour timepoint, middle row being the 4 hour timepoint and the last row being the 6 hour timepoint.

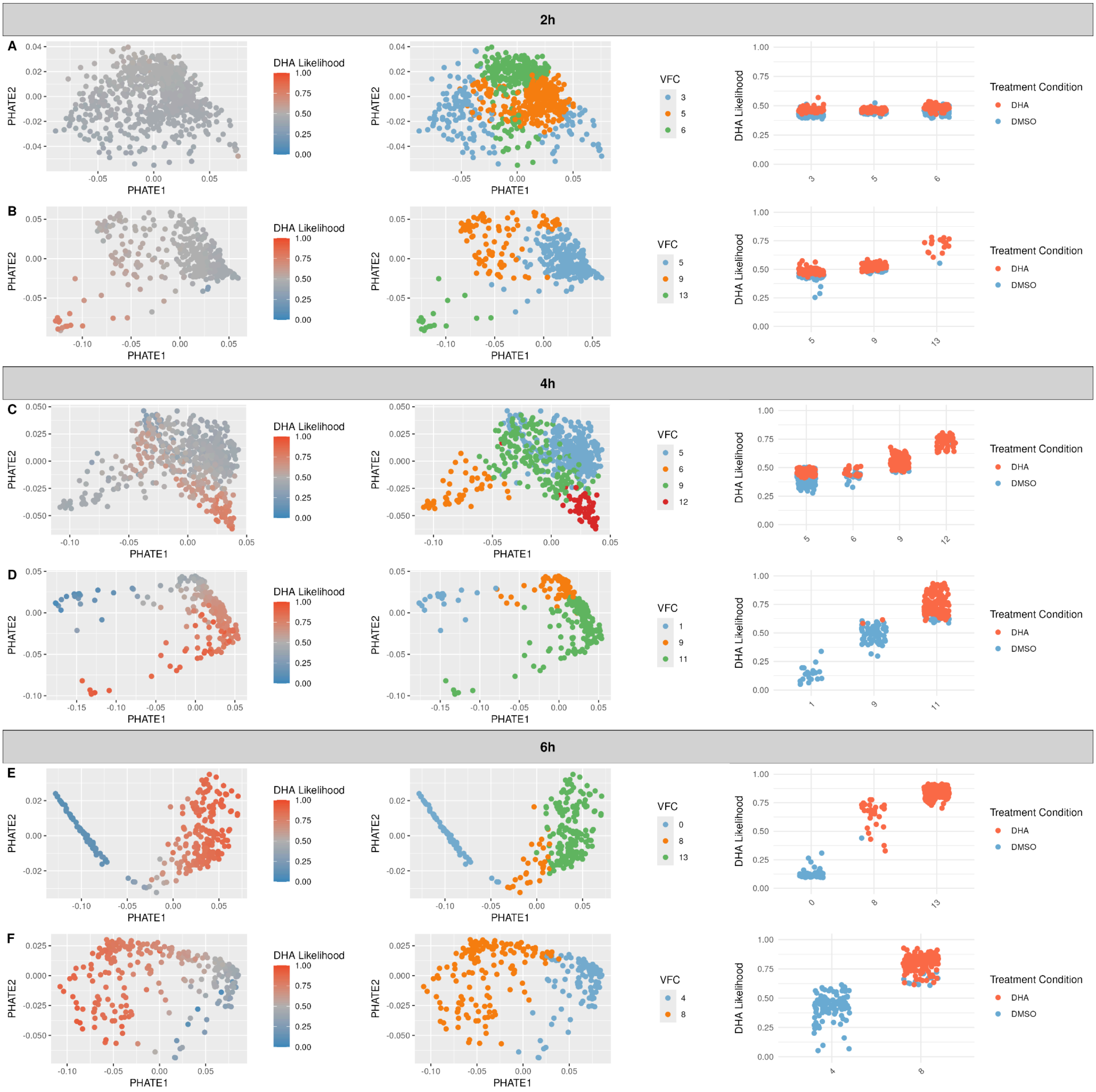


**Supplemental Figure S18. Vertex frequency clustering (VFC) results for sexually committed rings.** For each timepoint, the sexually committed ring stage was sub-clustered using Vertex Frequency Clustering. The first row (A,C,E) of each timepoint is K13$^{C580,}$ while the second row (B,D,F) is K13$^{580Y}$. For each timepoint, the clusters with the highest DHA likelihood were selected for differential expression.

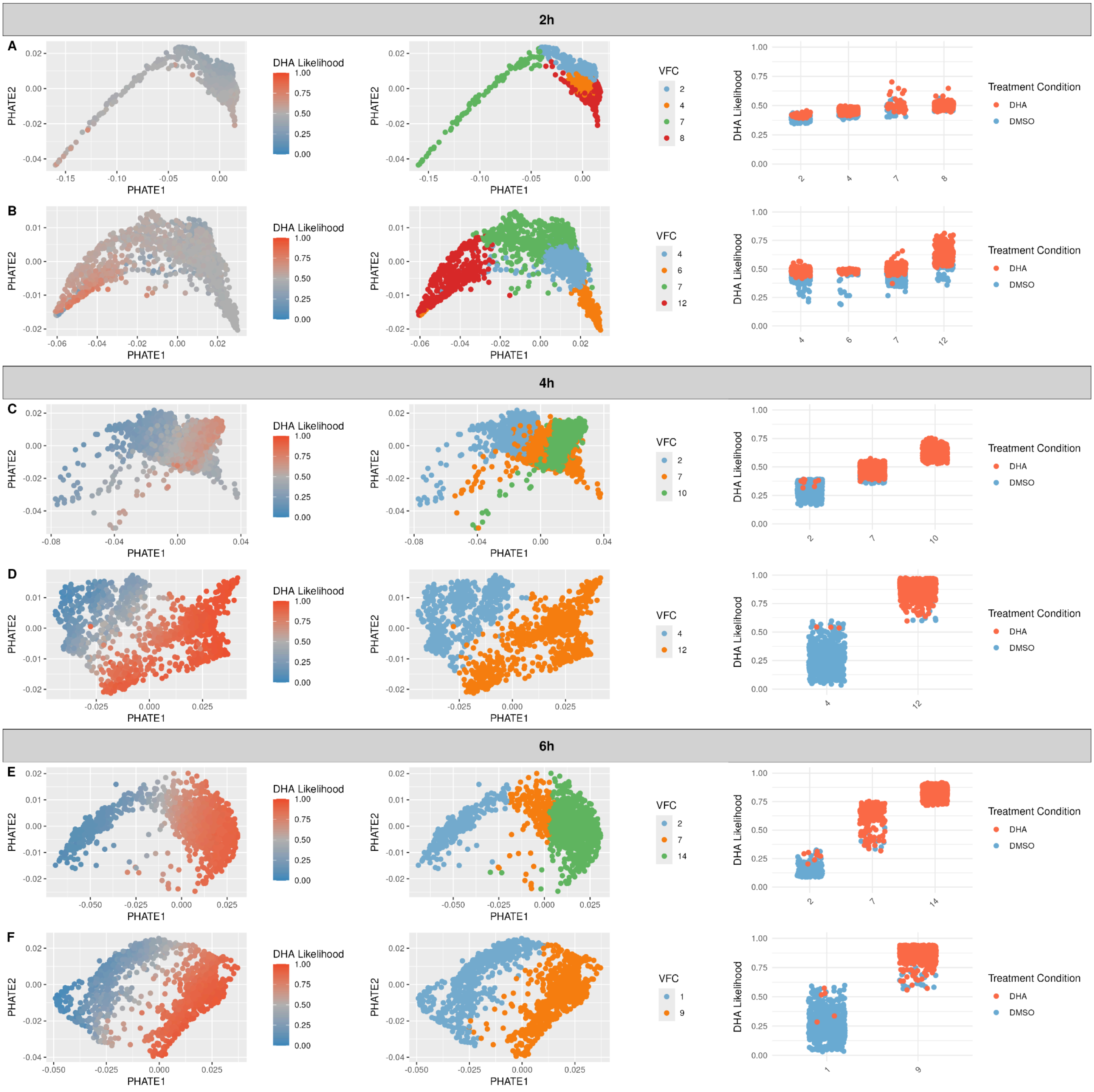


**Supplemental Figure S19. Vertex frequency clustering (VFC) results for rings.** For each timepoint, the ring stage was sub-clustered using Vertex Frequency Clustering. The first row (A,C,E) of each timepoint is $K13^{C580}$, while the second row (B,D,F) is $K13^{580Y}$. For each timepoint, the clusters with the highest DHA likelihood were selected for differential expression.

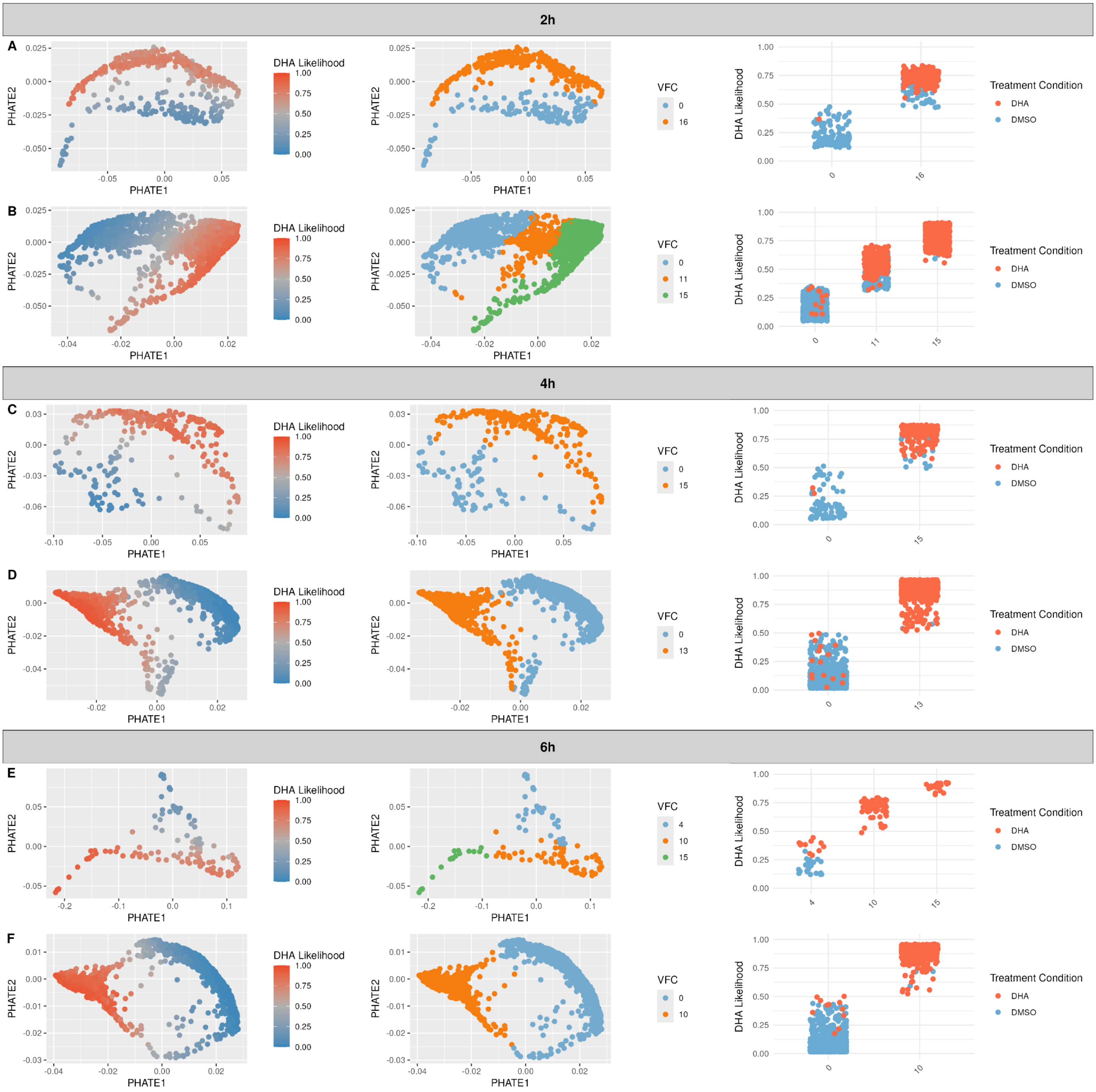


**Supplemental Figure S20. Vertex frequency clustering (VFC) results for trophozoites.** For each timepoint, the trophozoite stage was sub-clustered using Vertex Frequency Clustering. The first row (A,C,E) of each timepoint is K13$^{C580,}$ while the second row (B,D,F) is K13$^{580Y}$. For each timepoint, the clusters with the highest DHA likelihood were selected for differential expression.

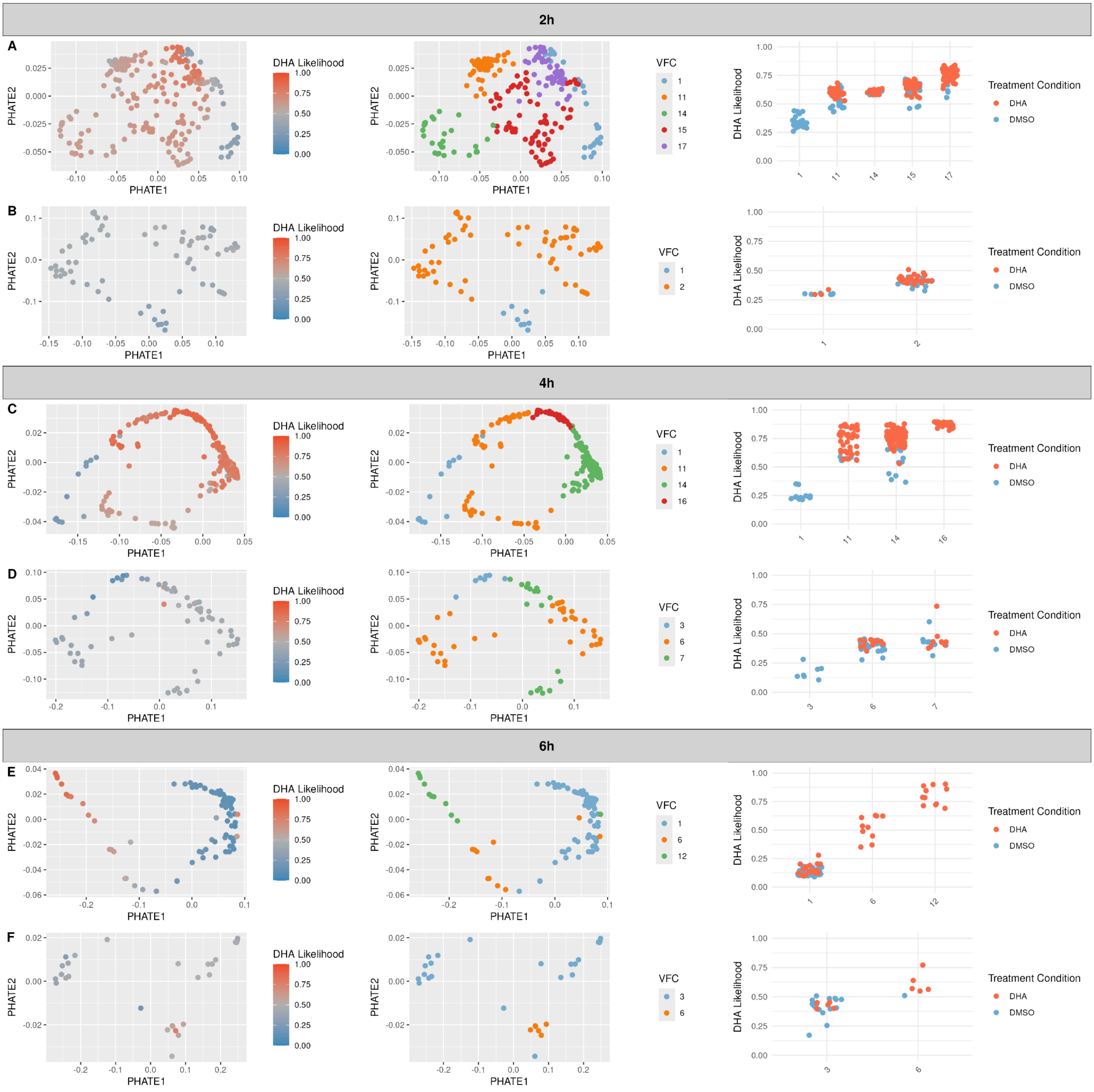


**Supplemental Figure S21. Vertex frequency clustering (VFC) results for schizonts.** For each timepoint, the schizont stage was sub-clustered using Vertex Frequency Clustering. The first row (A,C,E) of each timepoint is $K13^{C580,}$ while the second row (B,D,F) is $K13^{580Y}$. For each timepoint, the clusters with the highest DHA likelihood were selected for differential expression.

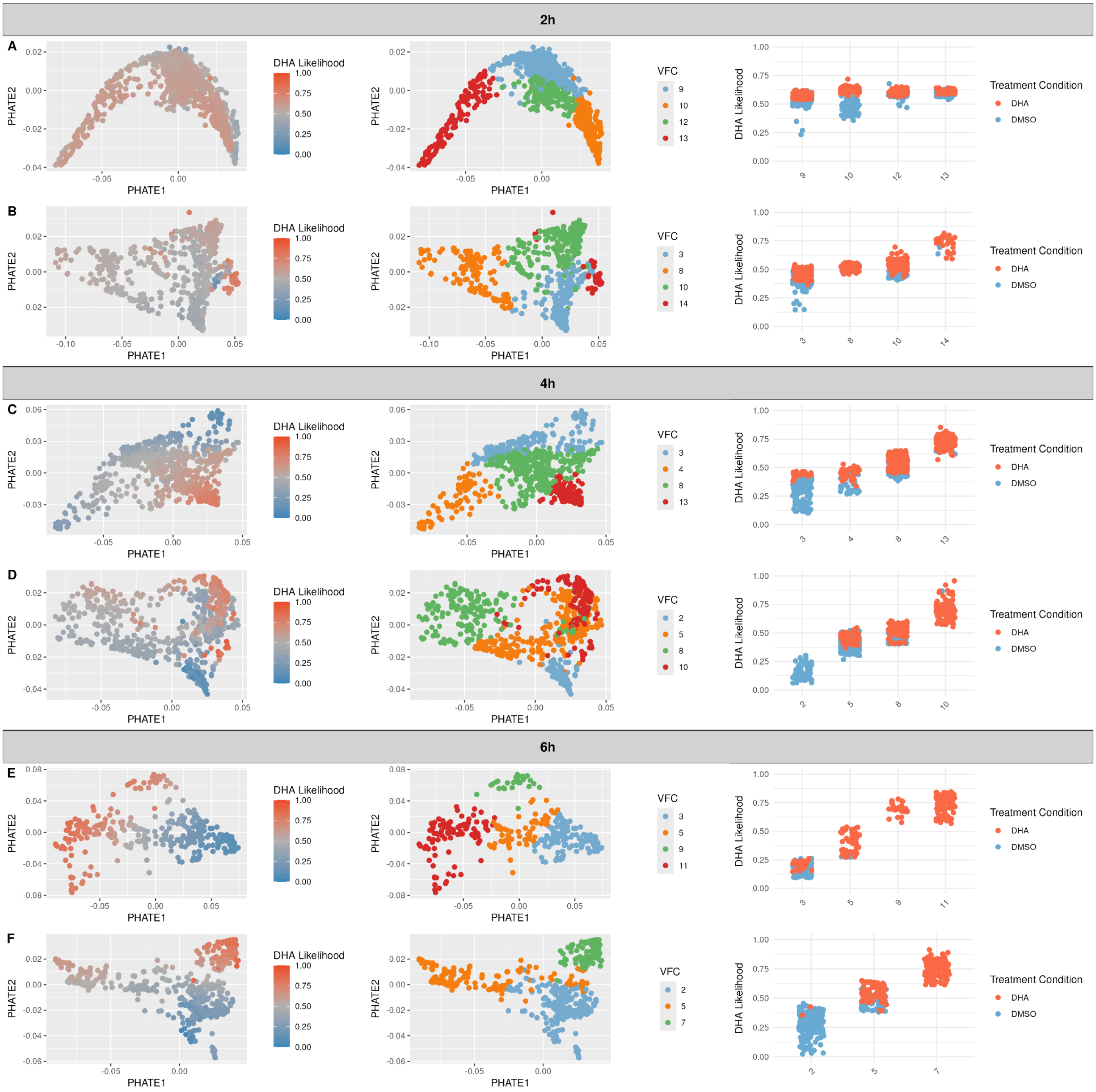


**Supplemental Figure S22. Vertex frequency clustering (VFC) results for gametocytes.** For each timepoint, the gametocyte stage was sub-clustered using Vertex Frequency Clustering. The first row (A,C,E) of each timepoint is K13$^{C580,}$ while the second row (B,D,F) is K13$^{580Y}$. For each timepoint, the clusters with the highest DHA likelihood were selected for differential expression.

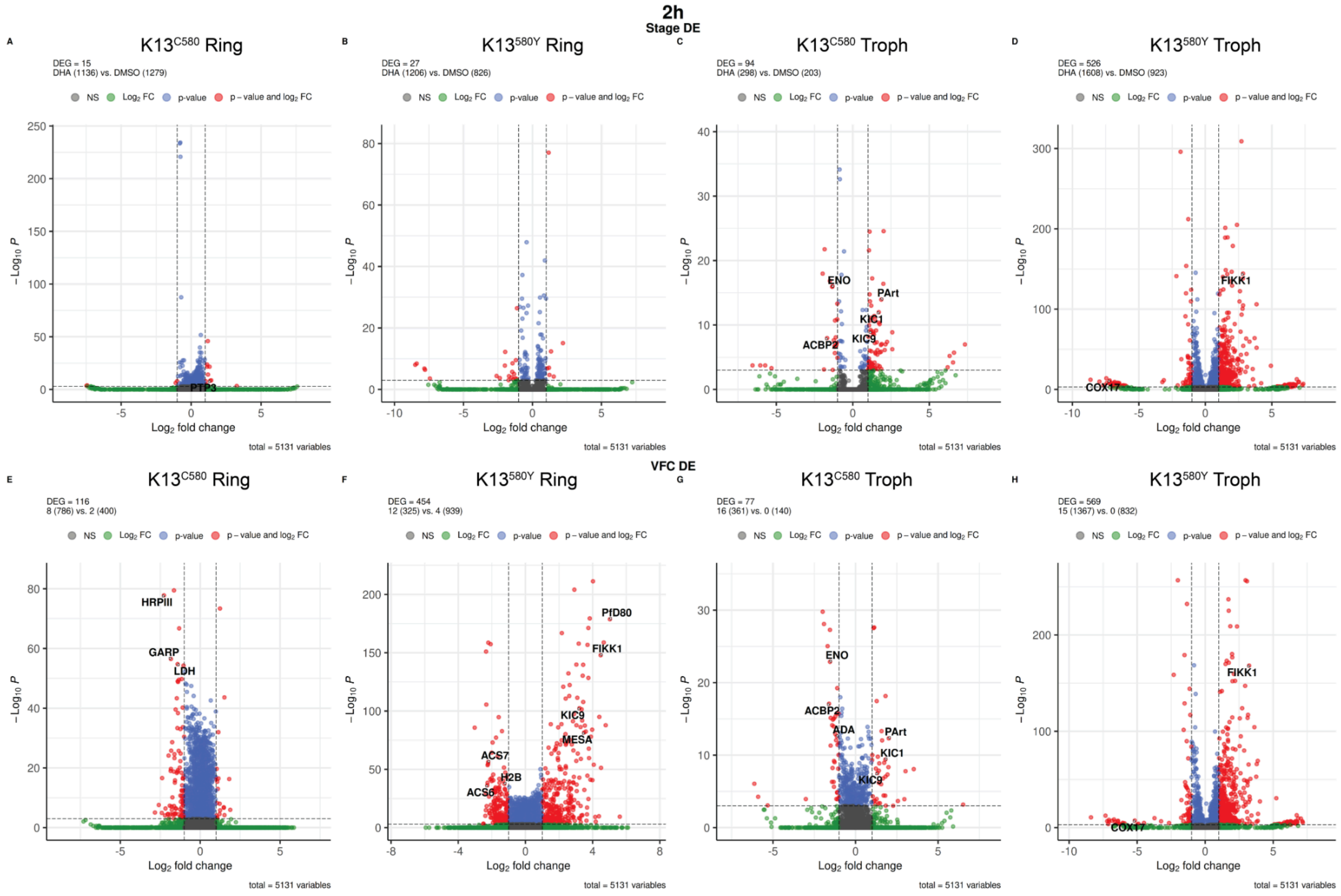


**Supplemental Figure S23. Similar pattern of differentially expressed genes in ring stages using VFC cluster differential expression compared to stage differential expression at the 2 hour timepoint.** Volcano plots in each panel show the average log2FC on the x axis and the negative log10 of the BH-adjusted p value output from MAST (17) on the y axis. Colors denote different levels of significance; non significant genes are in grey, genes with an absolute average log2FC > 1 are green, genes with a BH-adjusted p value < 0.001 are in blue and significant genes with an absolute log2FC > 1 and a BH-adjusted p value < 0.001 are in red. The first row represents differential expression results when comparing between DHA vs. DMSO conditions within a stage for the 2 hour timepoint, between K13$^{C580}$ rings (A), K13$^{580Y}$ rings (B), K13$^{C580}$ trophozoites (C) and K13$^{580Y}$ trophozoites (D). The second row represents differential expression results when comparing the VFC cluster with the highest DHA likelihood with the VFC cluster with the lowest DHA likelihood, between K13$^{C580}$ rings (E), K13$^{580Y}$ rings (F), K13$^{C580}$ trophozoites (G) and K13$^{580Y}$ trophozoites (H).

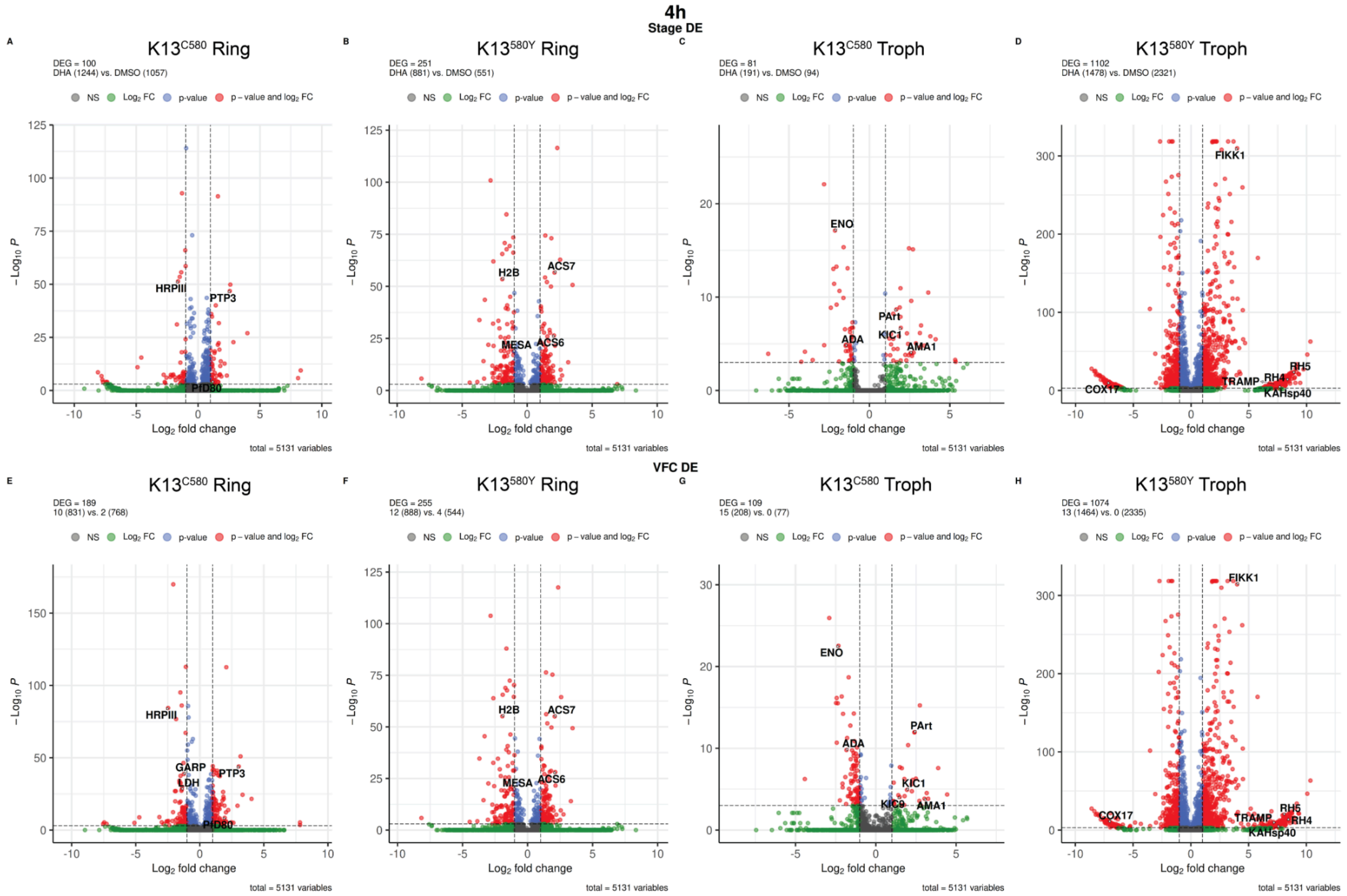


**Figure S24. Similar pattern of differentially expressed genes in ring stages using VFC cluster differential expression compared to stage differential expression at the 4 hour timepoint.** Volcano plots in each panel show the average log2FC on the x axis and the negative log10 of the BH-adjusted p value output from MAST (17) on the y axis. Colors denote different levels of significance; non significant genes are in grey, genes with an absolute average log2FC > 1 are green, genes with a BH-adjusted p value < 0.001 are in blue and significant genes with an absolute log2FC > 1 and a BH-adjusted p value < 0.001 are in red. The first row represents differential expression results when comparing between DHA vs. DMSO conditions within a stage for the 4 hour timepoint, between K13$^{C580}$ rings (A), K13$^{580Y}$ rings (B), K13$^{C580}$ trophozoites (C) and K13$^{580Y}$ trophozoites (D). The second row represents differential expression results when comparing the VFC cluster with the highest DHA likelihood with the VFC cluster with the lowest DHA likelihood, between K13$^{C580}$ rings (E), K13$^{580Y}$ rings (F), K13$^{C580}$ trophozoites (G) and K13$^{580Y}$ trophozoites (H).

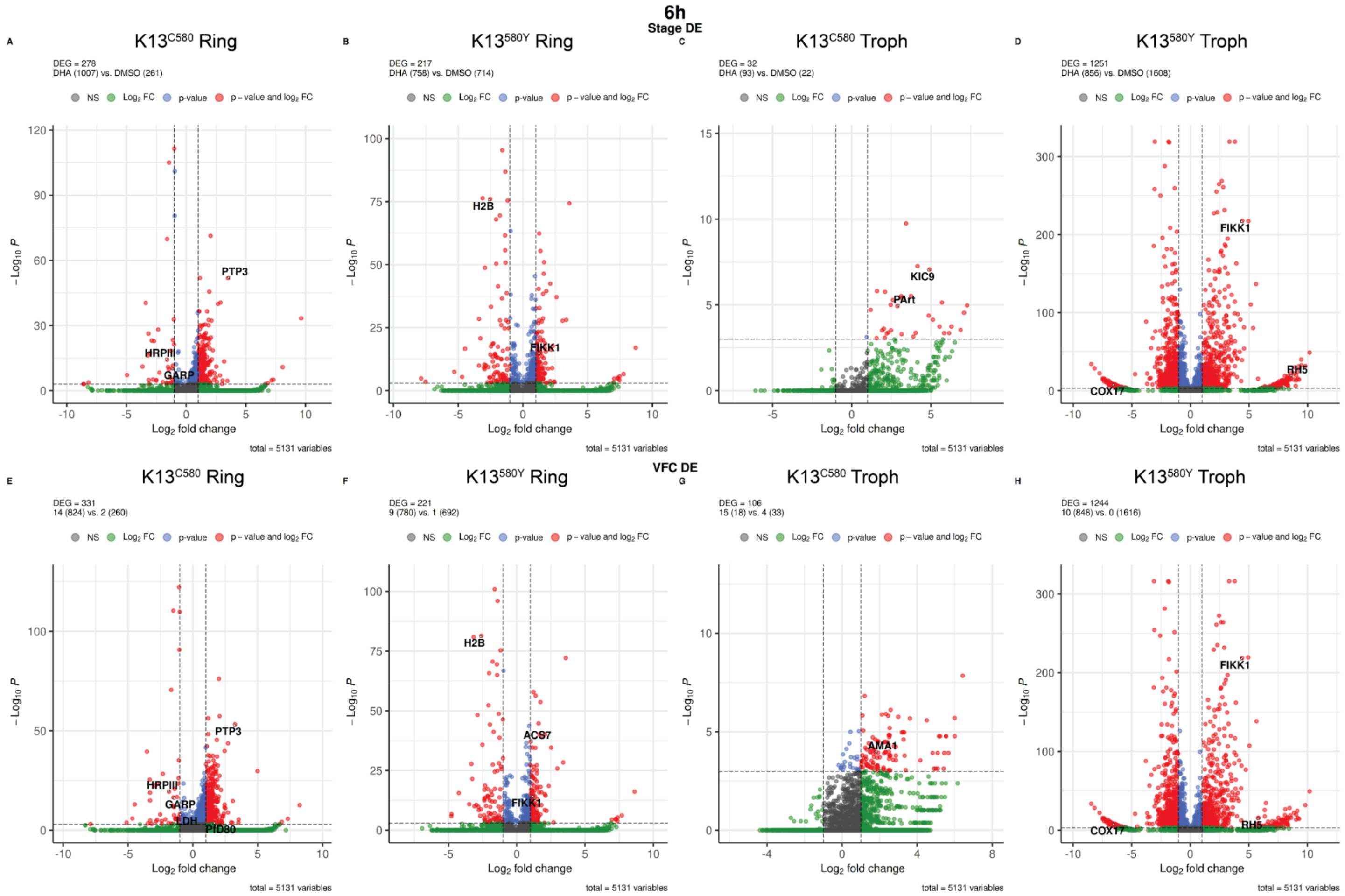


**Figure S25. Similar pattern of differentially expressed genes in ring stages using VFC cluster differential expression compared to stage differential expression at the 6 hour timepoint.** Volcano plots in each panel show the average log2FC on the x axis and the negative log10 of the BH-adjusted p value output from MAST (17) on the y axis. Colors denote different levels of significance; non significant genes are in grey, genes with an absolute average log2FC > 1 are green, genes with a BH-adjusted p value < 0.001 are in blue and significant genes with an absolute log2FC > 1 and a BH-adjusted p value < 0.001 are in red. The first row represents differential expression results when comparing between DHA vs. DMSO conditions within a stage for the 6 hour timepoint, between K13$^{C580}$ rings (A), K13$^{580Y}$ rings (B), K13$^{C580}$ trophozoites (C) and K13$^{580Y}$ trophozoites (D). The second row represents differential expression results when comparing the VFC cluster with the highest DHA likelihood with the VFC cluster with the lowest DHA likelihood, between K13$^{C580}$ rings (E), K13$^{580Y}$ rings (F), K13$^{C580}$ trophozoites (G) and K13$^{580Y}$ trophozoites (H).

**Supplemental Table S1 - Differential Expression Results over All Cell Stages**

See the uploaded Excel file.

**Supplemental Table S2 - Gene Ontology Enrichment over All Cell Stages, By Category**

See the uploaded Excel file.

**Supplemental Table S3 - Gene Ontology Enrichment over All Cell Stages, By Gene**

See the uploaded Excel file.

**Supplemental Table S4. Genotypes of Targeted Drug Resistance Marker Loci Captured with Molecular Inversion Probes.**

See the uploaded Excel file.
